# Revealing Notch-dependencies in synaptic targets associated with Alzheimer’s disease

**DOI:** 10.1101/2021.03.22.436438

**Authors:** A. Perna, S. Marathe, R. Dreos, L. Falquet, H. Akarsu, L. Alberi Auber

**Affiliations:** Department of Medicine, Unit of Pathology, University of Fribourg, Fribourg, Switzerland; Centre for Neuroscience, Indian Institute of Science, Bangalore, India; Swiss Institute of Bioinformatics (SIB), Lausanne, Switzerland; Biochemistry Unit, University of Fribourg and Swiss Institute of Bioinformatics, Fribourg, Switzerland; Swiss Integrative Center for Human Health, Fribourg, Switzerland

**Keywords:** Alzheimer’s disease, hippocampus, Notch, cell signaling, transcription, plasticity

## Abstract

Alzheimer’s disease (AD) is a progressive neurodegenerative disorder and the major cause of dementia. There is evidence that synaptic dysfunction and perturbation of Excitatory/Inhibitory (E/I) balance arise at the early stages of AD, altering the normal neural network activity, and leading to cognitive decline. Recent studies have identified Notch signaling as a contributor of neurodegenerative advancement including AD pathophysiology. As part of the efforts to understand molecular mechanisms and players involved in cognitive decline, we employed transgenic mouse models with Notch1 and RBPJK loss of function (LOF) in pyramidal neurons of the CA fields. Using bulk RNAseq. We have investigated the differential expression of Notch-dependent genes either upon environmental enrichment (EE) or upon Kainate injury (KA). We found a substantial genetic diversity in absence of both Notch1 receptor or Rbpjk transcriptional activator. Among differentially expressed genes, we observed a significant upregulation of Gabra2a in both knockout models, suggesting a role for Notch signaling in the modulation of E/I balance. Upon neuroexcitotoxic stimulation, loss of Rbpjk results in decreased expression of synaptic proteins with neuroprotective effects. We confirmed Nptx2, Npy, Pdch8, TncC as direct Notch1/Rbpjk targets and Bdnf and Scg2 as indirect targets. Finally, we translate these findings into human entorhinal cortex containing the hippocampal region from Alzheimer’s Disease patients performing targeted transcripts analysis. We observe an increased trend for Rbpjk and the ligand DNER but not Notch1 expression. On the other hand, neuron-specific targets, Nptx2, Npy, BDNF and Gabra2a are upregulated during the mild-moderate stage, and decline in the severe phase of the disease. These findings identify Notch as a promising signaling cascade to fine-tune in order to ameliorate synaptic transmission and memory deficits that occur during early phase of the Alzheimer’s Disease.

**Highlights:**

- Loss of canonical and/or non-canonical Notch1 signaling in pyramidal neurons of the hippocampal CA field mainly affects the post-synaptic compartment.
- In both RBPJKcKO and Notch1cKO mouse models there is upregulation of GABAergic receptor subunit alpha2 (Gabra2a).
- The plasticity genes: Npy, Nptx2,Pcdh8 and TncC with neuroprotective functions and known association with Alzheimer’s Disease are direct Notch/Rbpjk targets.
- During the mild-moderate stage of AD dementia, Notch canonical signaling promotes the expression of neuroprotective proteins, in the attempt of mitigating the effect of the excitatory-inhibitory imbalance. This activity is not observed during severe stages of the disease.

## 1. Introduction

Alzheimer’s disease (AD) is the most frequent form of age-related dementia[1]. It is a neurodegenerative disorder, clinically characterized by initial progressive memory deficits and eventually accompanied by a more global cognitive and intellectual decline, leading to severe dementia. Core neuropathologies in AD include the accumulation of the protein amyloidbeta (Aβ) and the development of neurofibrillary tangles (NFT), which have been associated with neuronal degeneration and clinical symptoms of dementia[2]. The AD brain is also characterized by extensive neuronal and synaptic loss in areas of the brain essential for cognitive and memory functions, such as cerebral cortex and hippocampus, often accompanied by astrogliosis[3] and microglial cell proliferation[4]. Additionally, pathogenic mechanisms including oxidative stress[5], mitochondrial changes[6] and cell cycle re-entry[7] occur already in the preclinical phase of dementia before the onset of cognitive symptoms [8, 9]. Beyond the typical hallmarks of AD, synaptic dysfunction is also of major importance. In particular, changes in excitatory and inhibitory synapses releasing glutamate and GABA, respectively, have been ascribed as early contributing factors to AD pathology[10]. Hippocampal hyperactivation and impaired deactivation of the default-mode network during memory-encoding have been observed in people with genetic risk for AD[11, 12], cognitively normal individuals with evidence for Aβ accumulation[13, 14] and people with early AD[15, 16]. Increase of oxidative or metabolic stress can also lead to excessive glutamatergic tone, which is thought to lead to neuronal loss in AD[17]. Furthermore, there is great evidence that during early stage of the disease soluble Aβ oligomers and amyloid plaques alter the function of local neuronal circuits and large-scale networks by disrupting the balance of synaptic excitation and inhibition (E/I balance) in the brain[18, 19, 20, 21]. Signaling cascades respond to synaptic activity and contribute to network plasticity through transcriptional and post-transcriptional regulation, influencing the biological repertoire of both neurons and glia. These adaptive changes are fundamental in supporting brain functions, such as learning and memory[22], but can also propagate neuropathological signals contributing to progressive neurological deficits seen in Alzheimer’s disease[23]. Notch1 signaling, an evolutionarily conserved pathway fundamental for brain development and neural stem cell maintenance [24], has been previously implicated in AD. In the adult brain, Notch1 is expressed in pyramidal neurons of the cortex[25] and hippocampus[26]. Prevalently postsynaptic, Notch1 regulates spine morphology, synaptic plasticity and memory processing[27]. While some of these functions can be attributed to the interaction of Notch1 with NMDA and Reelin receptors [26], this signaling cascade remains largely unexplored in the context of the adult brain.

Notch signaling can either be canonical or non-canonical. Canonical signaling is triggered by ligand-binding, resulting in the generation of Notch Intracellular Domain (NICD). This protein’s truncation associates with the transcriptional repressor RBPJK in the nucleus and leads to the recruitment of activator factors, turning RBPjK into an activator and creating a complex necessary to transcribe downstream targets. So far, only a few relevant target genes were found and widely accepted as canonical Notch targets[28]. Among others, Hes and Hey gene family members are confirmed direct targets of the NICD/RBPJK complex[29]. Non-canonical Notch signaling, instead, is independent of RBPJK and interacts with several pathways at either the cytoplasmic and/or nuclear level[30]. A few of the important cytoplasmic partners of Notch are the tyrosine kinase Abl, mTOR, PI3K, Wnt/β-Catenin, and NF-κB [31] as well as membrane receptors, ApoER and NMDAR [26]. It has been proposed that canonical signaling is involved in cell demise following injury (pathological condition), while non-canonical signaling plays a dominant role in synaptic plasticity (physiological condition)[31]. Notch1 imbalances have been described in Alzheimer’s disease: presenilins mutations, characteristic of familiar AD, can affect Notch1 processing/activation, given that amyloid-β peptide (Aβ) and NICD precursors are cleaved by the same protease, γ-secretase[32, 33]. On the other hand, an increased Notch1 expression was also reported in sporadic AD patients, suggesting a potential role for the signaling beyond γ-secretase dysfunction[34]. In *post-mortem* patients with sporadic AD condition Notch1 is profoundly delocalized and associated with both fibrillary tangles and amyloid plaques[35]. So far, It has been demonstrated that novel spatial navigation induces Notch activity in the neuronal ensemble of the hippocampus, regulating synaptic plasticity and memory formation[27]. On the other hand, hyperactivation of Notch signaling can cause neuronal demise following ischemic injury[36]. In AD brains, Notch1 signaling appears reduced in neurons suggesting that a loss of function in Notch signaling may preserve neuronal survival at the expenses of synaptic plasticity contributing to the manifestation of progressive memory loss in AD. Understanding the molecular cascade associated to Notch activation during neuropathological progression may help untangle the dense signaling network contributing to the worsening of AD. In the search for Notch downstream targets contributing to its function, we coupled a transcriptomic approach with *in silico* analysis and ChIP validation of discovered targets upon neuroexcitotoxicity. To narrow our analysis to neurons, We employed transgenic mouse models with neuron-specific deletion of either Notch1 receptor or RBPJK transcriptional activator to exclude glial contribution and explored the transcriptional program of the signaling either during physiological state (Short Enriched Environment, EE) or Kainic Acid (KA)-induced neuroexcitotoxicity, resembling the glutamatergic imbalance typical of early phase of AD[37]. Finally, we translated our findings into humans by performing targeted analysis of Notch components and target genes in entorhinal cortices of AD patients with moderate and severe dementia. We show that loss of canonical and/or non-canonical signaling in pyramidal neurons of the hippocampal CA field mainly affects the postsynaptic compartment and results in decreased glutamatergic transmission by indirect upregulation of GABAergic receptor subunit alpha2 (Gabra2a) reducing the excitatory tone. In contrast, we found decreased expression of synaptic proteins with neuroprotective effect, later confirmed to be under direct (Npy, Nptx2, Pcdh8) or indirect (BDNF, Scg2) control of Notch1/RBPJK. Collectively, these findings identify Notch as a promising signaling to finetune in order to ameliorate synaptic transmission and memory deficits that occur during early phase of the disease.

## 2. Materials and Methods

### 2.1. Animals

All experiments on mice were performed with permission of the local animal care committee, University of Fribourg (Protocol no. 2016_32_FR registered 01/01/2017). Male animals (2-4 months of age) were housed on a 12h light-dark cycle with access to food and water *ad libitum.* RBPJKcKO and WT littermate control (RBPJKflox/flox, and CamKII::Cre) mice were obtained by crossing RBPJKflox/flox mice[38] to the CamKII::Cre (T29-1) mouse line[39] on a C57BL6/129 background. Tissue specific Notch1cKO was obtained by crossing Notch1flox/flox mouse line[40] with neuron specific CamKII::Cre (T29-1) mouse line[39].

### 2.2. Human tissue

Frozen human entorhinal cortex tissue was generously provided by the Medical Research Council Brain Bank for Dementia Research, Oxford, UK, by Professor Thomas Montine from the Department of Pathology of Stanford Medicine and by the Netherlands Brain Bank (NBB). The use of human tissue has been approved by the Ethical Commission of the Brain Bank for Dementia UK (OBB443 registered 1/05/2017 and OB344 registered 1/02/2014), Stanford (Stanford IRB), and the Ethical Commission from the Canton of Fribourg and Vaud (N. 325/14). All experiments conducted on human tissue comply with the WMA Declaration of Helsinki. Individuals were divided into three groups according to *post mortem* pathological evaluation and Braak staging (Table 1).

**Table 1:** Summary of patients’ cohort

| Group | N. patients | F:M | Age (mean±SD) | Braak Stage (N.) |
| --- | --- | --- | --- | --- |
| CTL | 11 | 7:4 | 78±8.74 | I (4) - II (7) |
| MODERATE | 11 | 7:4 | 86±5.21 | III (3) - IV (8) |
| AD | 16 | 8:8 | 81±8.02 | V (6) - VI (10) |

Values are means ± standard deviations. CTL = Healthy Controls, Moderate = Moderate Alzheimer's Diseases, AD = Severe Alzheimer's Disease. F = female; M = male.

### 2.3. Treatments

To investigate the KA-mediated excitotoxicity, mice were single-housed for 30 min before the KA treatment. KA (Abcam, Cambridge, UK) was injected intraperitoneally (i.p.) at 25 mg/kg of body weight. This dosage was shown to induce seizures and neurodegeneration in the hippocampus[41]. Control mice were i.p. injected with an equivalent volume of vehicle (0.9% saline). Mice were observed for 2 h after KA treatment and returned to their home cage. In rodents, Short Enriched Environment (EE) mimics circumstances of a stimulating and interesting living environment that is conducive to learning and cognition[42]. To explore plasticity, mice were put in a cage with toys, nesting material, tubes, huts, and running wheels for one week. This environment provides sensory, cognitive and motor stimulation that are normally not part of standard animal housing.

### 2.4. Tissue processing

One week after environmental enrichment and 12 hours after Kainic Acid (KA) treatment, mice were transcardially perfused with 0.9% sterile saline solution, brains were harvested and cut into two hemispheres. One hemisphere was dissected to collect the hippocampus and further dissected in an ice-cold saline solution to obtain the Corpus Ammonis (CA) fields, removing the dentate gyrus[26]. The tissue samples were collected into Eppendorf and were flash-frozen in liquid nitrogen and stored at −80°C until further use. Another hemisphere was post-fixed in 4% PFA for 1 day, followed by immersion in 30% sucrose at 4°C sucrose and then embedded in an OCT block for cryosectioning at 35*μ*m thickness (Leica, Germany) and used for histological studies.

### 2.5. Immunohistochemistry

The mice sagittal tissue sections from the anti-freezing medium were taken and then mounted onto the superfrost glass slides, air-dried for 1 hour, and washed twice in distilled water 5 minutes each. To access epitopes slides were incubated at 65°C (water bath) for 20 minutes in 10mM Sodium citrate (pH 6) containing 0.05% Tween (Preheat buffer). Immunohistochemistry for GABAR2a (Novus Biologicals; NBP2-36560), Nptx2 (Abcam, ab69858) and GluR1 (Santa Cruz Biotechnology, sc13152) was conducted as previously [41]

### 2.6. RNA isolation and sequencing

Total RNA was extracted using peqGOLD TriFast reagent (Peqlab, Erlangen, Germany). Isolated RNA was quantified and the quality was assessed with a Nanodrop (NanoDrop2000, Thermo Scientific, Waltham, MA, USA). Synthesis and amplification of cDNA were performed as per the Illumina TruSeq RNA protocol. RNA integrity was determined with the Fragment Analyzer 5200 (Agilent). Samples with RNA integrity number (RIN) > 8 were used for the experiment. An input material of 1 *μg* of total RNA from each sample was used for library preparation. Illumina TrueSeq Combinatorial dual (CD) indexes were used during the ligation, DNA fragments enriched using the PCR to amplify the amount of DNA in the library. The quality of the libraries is determined using the Standard High sensitivity NGS Fragment analysis kit (DNF-474, 1-6000 base pair) on the Agilent Fragment analyzer (Agilent, USA), yielding approximately 260 bp size fragments. The cDNA libraries were pooled in equivalent amounts. The libraries were denatured and diluted using standard library quantification and quality control procedures recommended as per the NextSeqproto-col. For a sequencing control, PhiX library was prepared and combined with the pooled prepared libraries. A final concentration of 1.5 pM library was sequenced on Illumina HighSeq system to generate 20 million of 2 x75 bp paired-end reads per library.

### 2.7. Data Analysis

The raw RNA-seq data was first checked for its quality with FastQC. The rRNA contamination was reasonably low for all samples. Raw data were aligned against the mouse reference genome mm10 using TopHat2 and the count matrix was built with HTSeq package on the output of TopHat2, i.e the correctly mapped and paired reads. Approximately 80% of the reads mapped to the mouse genome. Thereafter, 17-32 million uniquely and correctly mapped reads remained for the differential analysis. Differential expression analysis was performed with DESeq2 using samples listed in Table 2 and comparison schemes explained in the experimental design. Clustering of the samples revealed that CR3 seemed to be closer to the KA than the other two CR samples. This suggests that the kainate-treatment did not lead to a differential expression or did not work as well in this animal. For this reason, the following analyses were performed without including CR3 in the design. The differential expression was reported in log2 fold changes (log2FC) and only significant differences (padj < 0.05) were taken into consideration. To minimize the detection of false positives, the Benjamini-Hochberg correction is used[43].

**Table 2:** Summary of mouse samples used for RNA-seq

| Sample | Description | Genetic Background | Treatment | Pool |
| --- | --- | --- | --- | --- |
| CC1 | Control cage replicate 1 | wild-type (C57BL/6J) | control | 3 |
| CC2 | Control cage replicate 2 | wild-type (C57BL/6J) | control | 3 |
| CC3 | Control cage replicate 3 | wild-type (C57BL/6J) | control | 3 |
| CN1 | Mutant replicate 1 | Notch1cKO | Short Enriched Environment | 1 |
| CN2 | Mutant replicate 2 | Notch1cKO | Short Enriched Environment | 1 |
| CN3 | Mutant replicate 3 | Notch1cKO | Short Enriched Environment | 1 |
| CR1 | Mutant replicate 1 | RBPJcKO | Kainate 12h | 1 |
| CR2 | Mutant replicate 2 | RBPJcKO | Kainate 12h | 1 |
| CR3 | Mutant replicate 3 | RBPJcKO | Kainate 12h | 1 |
| EE1 | Control short-EE replicate 1 | wild-type (C57BL/6J) | Short Enriched Environment | 3 |
| EE2 | Control short-EE replicate 2 | wild-type (C57BL/6J) | Short Enriched Environment | 3 |
| EE3 | Control short-EE replicate 3 | wild-type (C57BL/6J) | Short Enriched Environment | 3 |
| KA1 | Control kainate replicate 1 | wild-type (C57BL/6J) | kainate 12h | 3 |
| KA2 | Control kainate replicate 2 | wild-type (C57BL/6J) | kainate 12h | 3 |
| KA3 | Control kainate replicate 3 | wild-type (C57BL/6J) | kainate 12h | 3 |

CC = cage controls, EE = enriched environment, KA = kainic acid, CN = Notch1cKO exposed to EE, CR = RBPJcKO with KA treatment.

### 2.8. Motif enrichment analysis

The RPBJK motif matrix (PWM) was downloaded from Ho-comoco database[44] and mapped to the mouse genome using PWMscan[45] with a p-value of 1e-4. Motif occurrence analysis around promoters (2KB window around annotated transcription start sites) of up and down regulated genes was performed in R using the ‘rtracklayer’ package[46]. These were then compared to the number of motif occurrences in the same number of promoters randomly picked from the expressed genes and repeated 500 times to generate a strong confidence interval.

### 2.9. Gene Ontology

#### 2.9.1. GOrilla

Gene Ontology enrichment analysis was performed with GOrilla tool[47] (http://cbl-gorilla.cs.technion.ac.il/), using two unranked gene lists (target and background). Target lists were generated for both up- and down-regulated genes belonging to the conditions CR versus KA and CN versus EE. For Kainic acid treatment, we chose to use differential expressed genes from the KA versus CC comparison as background list in order to spot just GO enrichment terms relative to the lack of RBPJK protein.

#### 2.9.2. REVIGO

REViGO Bubbleplots[48] of the Enriched GO Cluster representative of up- and down-regulated genes in RBPJKcKO and Notch1cKO were generated using GOrilla terms from the enrichment analysis mentioned before. The bubbleplot shows the cluster representatives (i.e. terms remaining after the redundancy reduction) in a two dimensional space derived by applying multidimensional scaling to a matrix of the GO terms’ semantic similarities[49]. Bubble color indicates the p-value; size indicates the frequency of the GO term in the database (bubbles of more general terms are larger).

#### 2.9.3. SYNGO

SynGO is a public knowledge-based web-tool that provides ontologies for the synapse. It relies on annotations based solely on published experimental evidence and curated by world-leading experts in synapse biology[50]. We performed the analysis for the CR versus KA and CN versus EE conditions using SynGO webtools (https://www.syngoportal.org/geneset.html). We looked at the gene count within the Biological Process (BP) and Cellular Component (CC) categories and we showed the analysis results using sunburst plots.

### 2.10. Gene Set Enrichment Analysis (GSEA)

Gene Set Enrichment Analysis[51] (https://www.gsea-msigdb.org/gsea/index.jsp) is a robust computational method that determines whether an *a priori* defined set of genes shows statistically significant, concordant differences upon KA treatment and RBPJK knockout. The analysis was performed for KA versus CC and CR versus KA datasets. Gene sets are available from Molecular Signatures DataBase (MSigDB, https://www.gsea-msigdb.org/gsea/msigdb/index.jsp) and for this analysis the HALLMARK collection of gene sets was used. Briefly, GSEA calculates an enrichment score (ES) that reflects the degree to which a gene set is overrepresented at the extremes (top or bottom) of the entire ranked list of differential expressed genes - where genes are ranked according to the expression difference (signal/noise ratio). The ES is calculated by walking down the list, increasing a running-sum statistic when it encounters a gene that is in the gene set and decreasing it when it encounters genes that are not. The enrichment score is the maximum deviation from zero encountered in the random walk; it corresponds to a weighted Kolmogorov-Smirnov-like statistic. The software then estimates the statistical significance (nominal P value) of the ES and, when an entire database of gene sets is evaluated, GSEA adjusts the estimated significance level to account for multiple hypothesis testing. GSEA first normalizes the ES for each gene set to account for the size of the set, yielding a normalized enrichment score (NES). It then controls the proportion of false positives by calculating the false discovery rate (FDR) corresponding to each NES. GSEA was run according to default parameters: collapses each probe set into a single gene vector (identified by its HUGO gene symbol), permutation number = 1000, and permutation type = “gene-sets”. In order to better visualize GSEA results from both analysis, we used Enrichment Map[52] (Cytoscape plugin) and to further analyze relationships between enriched genes in a given genes set, we used GeneMania[53] that finds genes likely to share function based on their interactions.

### 2.11. Chromatin Immunoprecipitation (ChIP)

For ChIP experiment we used the previously TF-ChIP protocol [54]. Briefly, after tissue homogenization, DNA and proteins were reversibly crosslinked with short formaldehyde incubation (1% FA-PBS) to maintain the association of proteins with their target DNA sequence. Cells were lysed with ionic lysis buffer and chromatin fragmented to 300 bp using ultrasounds. The lysate was first precleaned by incubation with sepharose beads and once nonspecific DNA fragments were removed, it was incubated overnight with anti-RBPJK (Cell Signaling; #5313) and anti-NICD (Cell Signaling; #4147). In the negative control sample, Fetal Bovine Serum (FBS) was added. Antibody-protein-DNA complexes were then precipitated using sepharose beads. Washes and elution enriched for Notch signaling target genes that once purified were ready for PCR.

### 2.12. RNA extraction and RT-PCR

For both mouse and human samples, total RNA was extracted using peqGOLD TriFast reagent (Peqlab, Erlangen, Germany) and quantified using Qubit 3.0 Fluorometer (High sensitivity, Invitrogen). RNA integrity was assessed by capillary electrophoresis using Fragment Analyzer (Advanced Analytical AATI, United States) and just samples with RQN > 5 were selected. For RNA retrotranscription and RT-PCR it was used GoTaq 2-Step RT qPCR System (Promega) according to the manufacturer’s instructions and it was performed using Mic qPCR Cycler (BioMolecular Systems, USA). Expression levels of genes of interest were determined using the CT method; the levels of the mRNAs of interest were normalized against the levels of the housekeeping gene, β-actin (mouse samples) and RPL13A (human samples). Correlation analysis (Spearman’s Rank) was conducted using R and significance tested using Student’s t-test.

### 2.13. PCR Primers

Primer pairs for RNAseq validation, ChIP assay and human RT-PCR are listed in Table 3. All primers were obtained from Microsynth, Switzerland.

**Table 3:**
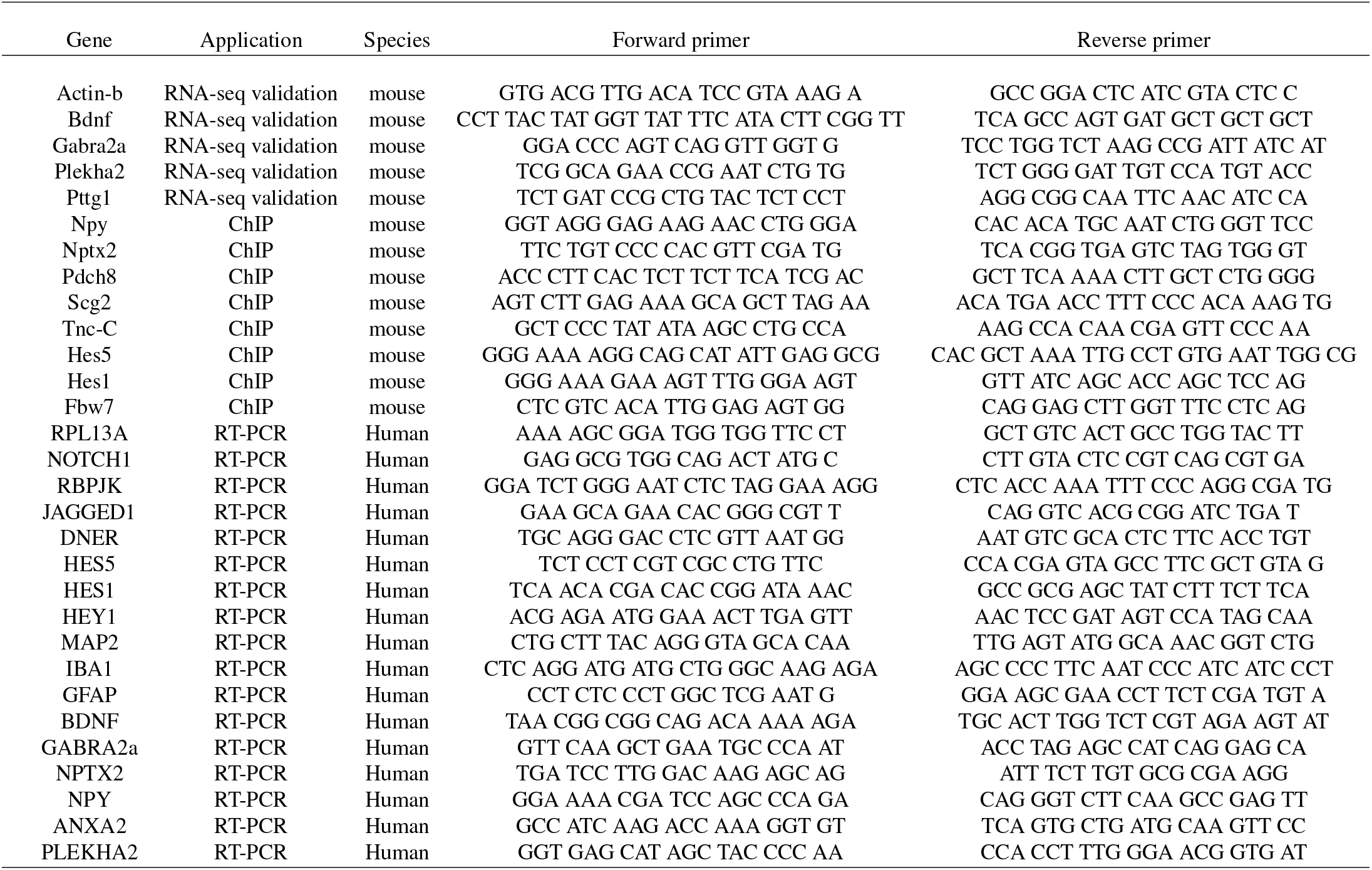
List of primers

## 3. Results

### 3.1. Experimental design for Notch targets discovery in neurons

To narrow down the understanding of Notch signaling in neurons we employed the Notch1cKO and RBPJKcKO transgenic mouse models, which lack the expression of respectively Notch1 or RBPJK in the hippocampal CA field (CamKII2-T291:cre) and allow to identify Notch-dependent targets in neurons excluding any other brain cell type contribution where Notch signaling is also represented. The Notch1cKO mouse model has been previously employed by our group to study the involvement of Notch1 in neuronal plasticity and memory/learning processes [27,26] and is useful in the identification of either direct canonical or non-canonical targets of the Notch1 receptor. The RBPJKcKO mouse model has been used to study the contribution of Notch canonical signaling to neurodegeneration and it is used to identify putative canonical targets of Notch1 in the context of neurons [41]. The previous evidence supported that Notch operates in a dual role with non-canonical signaling prevailing during physiological conditions and canonical signaling overriding upon pathological stimuli. For this reason we investigated the two transgenic models, RBPJKcKO (CR) and Notch1cKO (CN) under Kainate injury (KA) and environmental enrichment (EE), respectively. Only CA fields from all mice were recovered to capture the effect of the targeted loss-of-function (LOF) in the CA region even when using whole RNAseq. The resulting differential expressed genes (DEGs) are interrogated based on GO analysis and direct targets are explored. The DEGs are validated using RT-PCR in the rodent model and transposed to human brains of AD patients with increasing severity of the disease as compared to controls (Figure 1).

**Figure 1:**
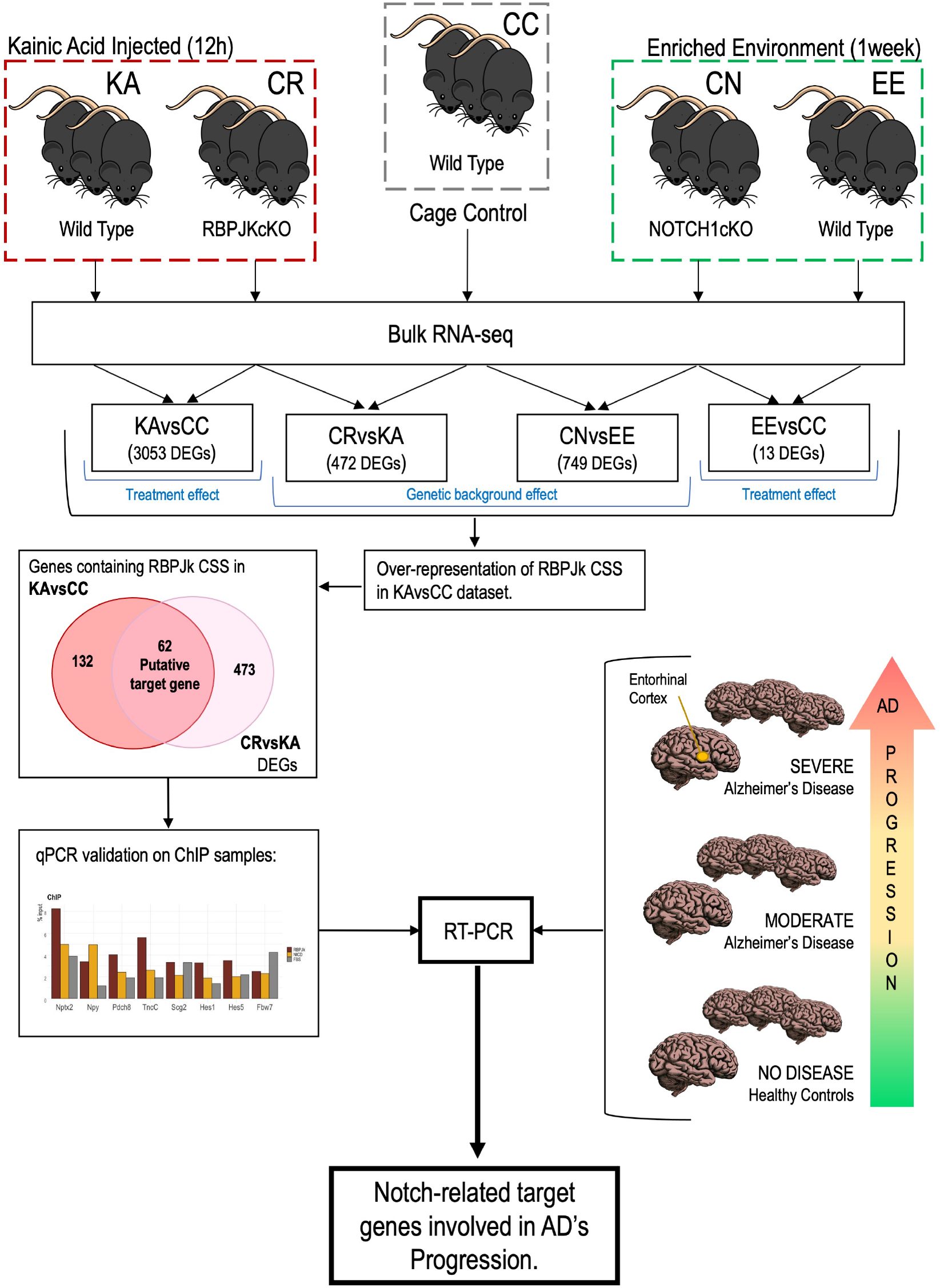
Experimental Design. Flowchart of study architecture and salient experimental steps aimed at identifying Notch/Rbpjk signaling targets in hippocampal ensembles. Wildtype cage control mice (CC), wildtype (EE) and Notch1cKO (CN) mice exposed to short enriched environment, wildtype (KA) and RBPJKcKO (CR) mice exposed to kainate injury;

### 3.2. RNAseq analysis of Notch-dependent targets

Using bulk RNAseq from CA dissected hippocampal regions, we have investigated the differential expression of Notch dependent genes either upon EE or KA. For data analysis, different comparison schemes were carried out (Figure 1): Comparison of the effect of EE or Kainate injury with normative conditions (CC) allows to determine treatment-dependent genetic changes. After 12 hours from induction induces a significant change in gene expression: 3053 genes are differentially expressed (DEGs), padj < 0.05 and 136 DEGs have a 2 Fold change cutoff (blue dots) with the majority of the genes in this subgroup being upregulated, 89% (Figure 2A). The synaptic gene ontology (SYN-GO) analysis of the up-regulated DEGs shows an enrichment in 7 postsynaptic genes of the postsynaptic density (Adra2c, Gsg1l, Ptprf, Sorcs3, Itga5, Slc6a8 and Nptx2) and 4 pre-synaptic genes regulating presynaptic vesicle sorting (Scg2, Rab26) and synaptic membrane (Nptx2 and Sv2c). Notch1 is also significantly but more modestly upregulated (Log2 Fold change= 0.56) to a similar extent as its canonical target Hes1 (Log2 Fold change= 0.66) and associated Notch-gene BDNF (Log2 Fold change= 0.69). While its other canonical target Hes5 appears down-regulated (Log2 Fold change= −0.89). In line with previous research, KA administration induces aberrant regulation of genes involved in neurotransmission like Secretogranin II (Scg2), shown to increase expression in rat hippocampus following KA intoxication[55], Neuropeptide Y, Npy and its receptors Npy1r and Npy2r, known to be involved in seizure modulation[56] and Huntingtin-associated protein 1 (Hap1).

**Figure 2:**
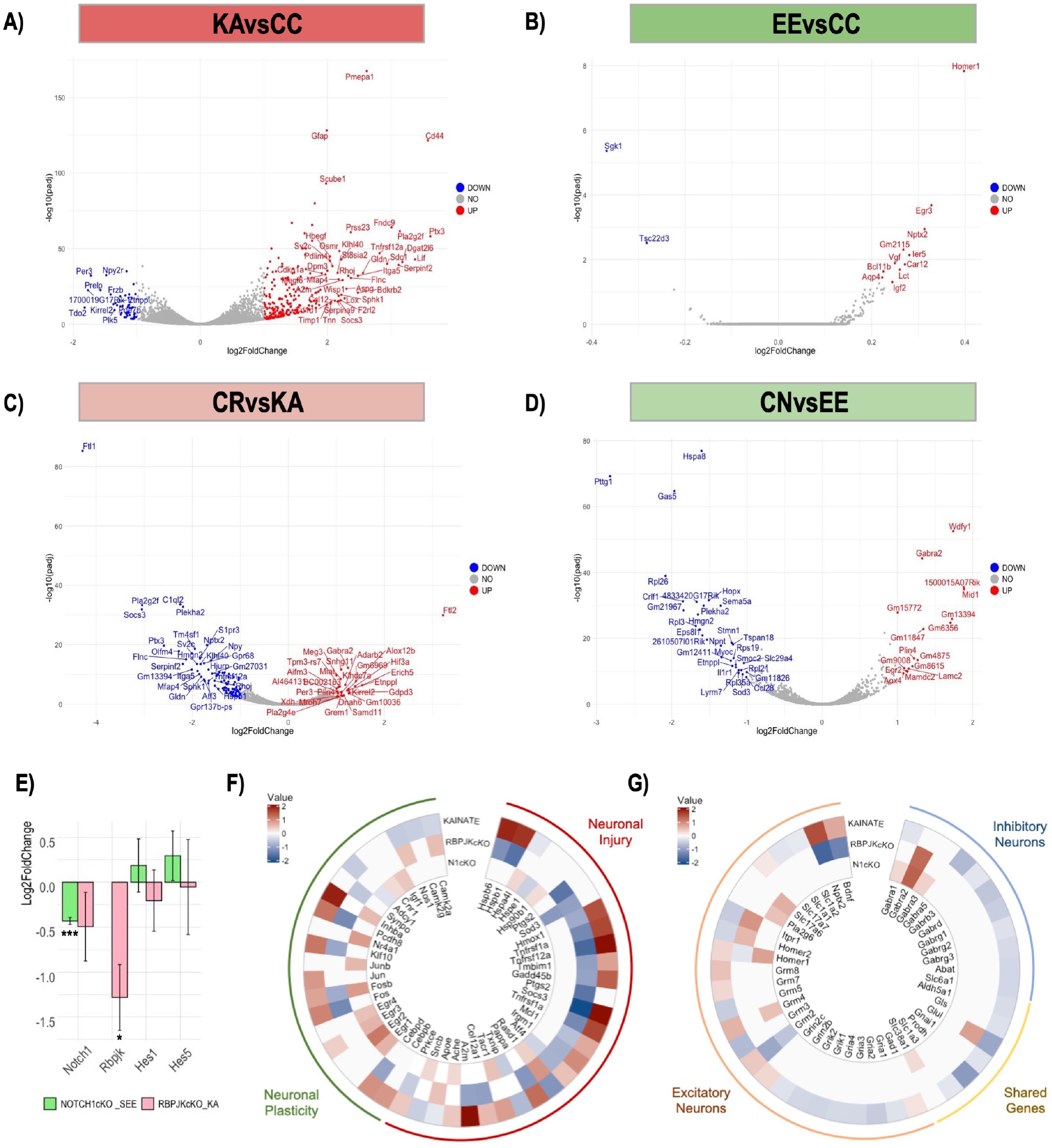
RNA sequencing analysis of mice lacking either RBPJK or Notch1 in hippocampal neurons. Volcano plots with log2-fold change (X-axis) and −log10 p-value (Y-axis) of differential expressed genes from comparison of **A)** Kainic Acid (KA) injected (25 mg/kg,12h) and wild type (CC) mice, **B)** mice exposed to an Enriched Environment (EE) for 1 week and wild type mice (CC), **C)** RBPJk conditional Knock-Out (CR) and wild type mice (KA), both Kainic Acid injected, **D)** Notch1 conditional Knock-Out (CN) and wild type mice (EE), both sacrificed 1 week after exposure to the enriched environment. **E)** RTPCR analysis of Notch1 signaling components in RBPJKcKO mice upon neuroexcitotixity as compared to KA and NOTCH1cKO mice under Short Enriched Environment as compared to EE. Circular heatmaps show differential expression (Log2FC) of genes involved in **F)**) neuronal plasticity/neuronal injury and **G)** excitatory/inhibitory neurons. *** indicates P values < 0.001 and * is for pvalue < 0,05.

We also observe robust upregulation of genes involved in reactive astrogliosis, like glial fibrillary acidic protein Gfap[57] and the proinflammatory adhesion molecule CD44[58]. Neuroexcitotoxicity induced also general upregulation of Notch signaling components, which is repressed by loss of RBPJK, supporting the involvement of Notch signaling in neuronal demise after excitotoxicity[41]. On the other hand, one week of EE causes only subtle changes (13 DEGs, padj < 0.05, with Log2 Fold changes between 0.2 and 0.4). 5 neuronal plasticity genes appear slightly upregulated (Ier5, Nptx2, Egr3, Homer1) and the rest of the DEGs belong to glio-vascular homoeostasis (Figure 2B). We have previously demonstrated that the loss of RBPJK in hippocampal neurons is neuroprotective and we have identified cell-cycle reentry as one of the putative neurodegenerative mechanisms regulated by Notch/RBPJK signaling [41]. To further expand our understanding of the gene targets contributed by the loss of canonical Notch signaling under Kainate stimulus, KA-injected RBPJKcKO (CR) mice were compared to KA mice after 12 hours of challenge. Of the 461 significant DEGs with a padj < 0.05, 116 DEGs show an equal or larger 2-fold cutoff with a concentration in downregulated hits (79%, Figure 2C). This is in opposition to large gene upregulation upon KA treatment (Figure 2A) and supports that Notch/RBPJK signaling is a transcriptional activator in condition of injury (Figure 2E). Statistical over-representation analysis of GOs using the REVIGO tools for cellular components on the upregulated genes, shows a localized enrichment for dendrites, neuronal membrane parts and GABAergic synapse while downregulated hits are more concentrated in the extracellular space besides conserving a synaptic localization (Supplementary Figure 1A). SYNGO analysis represented as sunburst chart illustrates the synaptic terms occurrence and confirms the synaptic identity of the differentially expressed hits with a concentration of post-synaptic genes (Supplementary Figure 1B) supporting the specificity of the RBPJKcKO model for studying Notch-dependent processes in neurons. Among differentially expressed genes, Npy, the synaptic protein Neuronal Pentraxin II (Nptx2), the lipid metabolism mediator Plekha2 and Bdnf neurotrophin which are upregulated after KA administration (Figure 2A) are repressed in absence of RBPJK (Figure 3B), raising the idea that these genes may be under either direct or indirect control of the signaling. Further elucidating the role of Notch1 in memory processing, we have investigated the contribution of Notch1 in condition of increased synaptic plasticity (CN) and compared the Notch1cKO to mice exposed to short environmental enrichment (EE). While the EE paradigm did not elicit substantial changes, the comparison shows 749 DEG with a padj < 0.05 and a smaller proportion of genes, 48, with a 2-fold change cutoff for the large part downregulated (65%), reflecting the Notch1 loss of function (Figure 2D). GO analysis using the REVIGO tools for cellular components on the upregulated genes, shows an enrichment for neuronal and synaptic components as well ribosomal subunits, while down-regulated genes are concentrated in the extracellular space and synapse (Supplementary Figure 1C). To confirm the topology of the gene hits, SYNGO reveals that also in case of Notch1 loss in neurons, the majority of the hits are localized post-synaptically and distributed between ribosomal genes and neurotransmitter transmembrane receptors (Supplementary Figure 1D). Notch1 LOF in hippocampal neurons reduced expression of Insulin degrading enzyme (Ide), the synaptic vesicle protein Synaptoporin (Synpr) and upregulated the synaptic anchoring Homer protein homolog 1 (Homer1) and Perilipin 4 (Plin4), coating protein of lipid droplets. Furthermore lack of Notch1 receptor in EE condition induces enrichment in GABAergic signaling through upregulation of Plekha2 and the GABAergic Receptor 2 (Gabra2), similarly to the loss of RBPJK (Figure 2C and 2D). To validate the ablation of Notch and RBPJK components in the respective conditional KO mice, we performed RT-PCR on whole hippocampal tissue (n= 4 for each genotype/condition) and confirmed that Notch1 is significantly but moderately reduced (30%, p<0.001) in Notch1cKO (Figure 2E) suggesting either a partial deletion or an increased expression of Notch1 in other cells than neurons as previously observed [27]. On the other hand RBPJK is significantly reduced (60%, p<0.05) in the RBJKcKO reflecting the loss of canonical signaling (Figure 2E). To narrow the understanding of the contribution of transcriptional versus default (canonical and non-canonical) Notch signaling, we have utilized literature-based gene pools for neuronal plasticity and injury. Overall the KA challenge induces the expression of the selected hits, while in absence of RBPJK directionality of expression is overall reversed with a conserved trend also in the Notch1cKO for the transcription factor Atf4 and Sod3 (Figure 2F). Further dissecting the contribution of Notch1 and RBPJK, selected gene-sets for excitatory neurons reveals a Notch-canonical repression of Nptx2, Bdnf and Grm2, accompanied by an activation of the inhibitory component, Gabra2a, also present in the Notch1cKO (Figure 2G). Overall, the absence of RBPJK reveals a more robust effect on gene expression both based on the ubiquitous deletion of Notch signaling and the KA treatment condition. Overall the RNA-seq analysis using the LOF model for Notch1 and RBPJK supports the bimodal role of Notch in memory formation and the neuronal response to excitotoxic insult.

**Figure 3:**
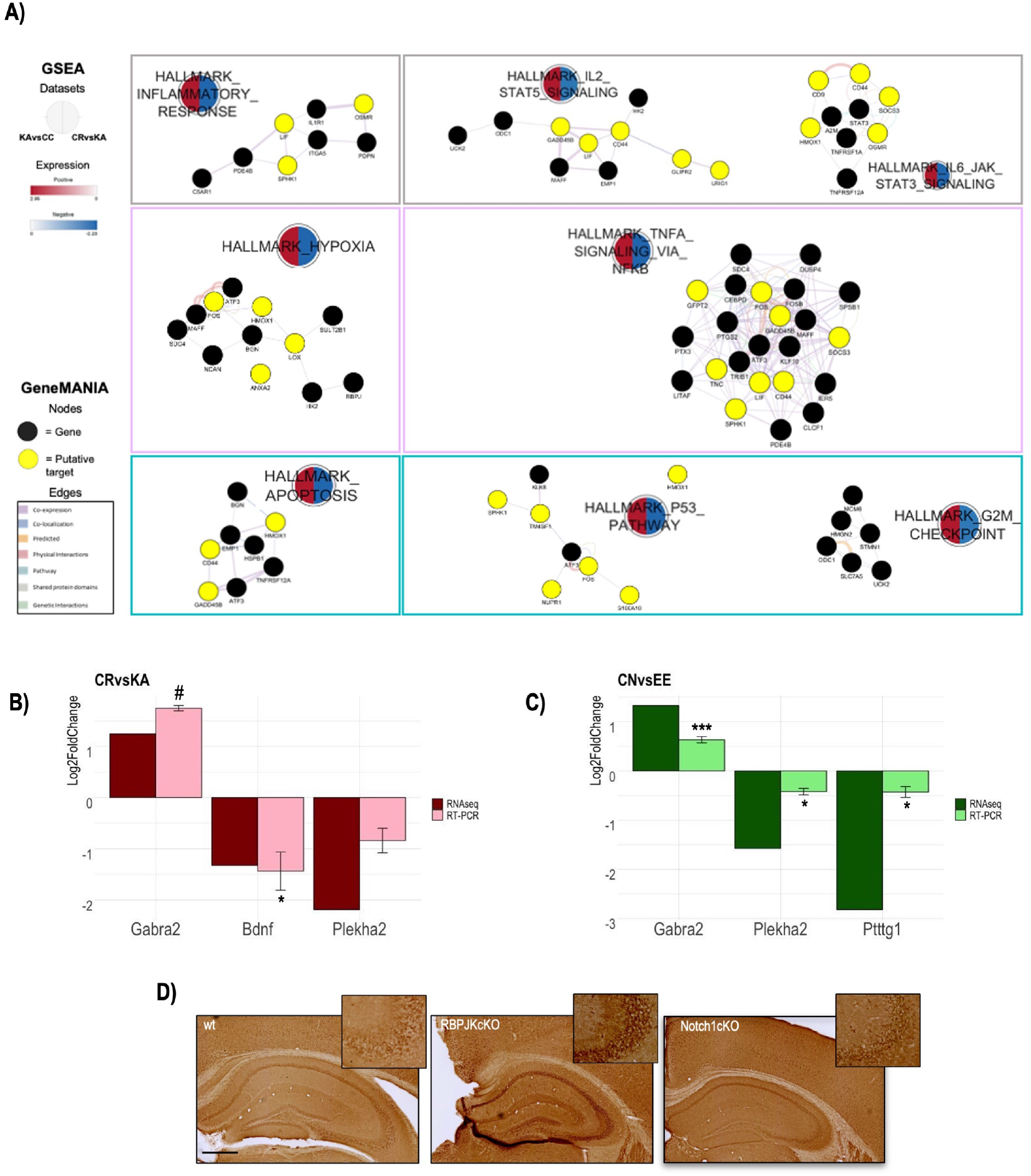
Transcriptional changes in Notch1cKO mice after SEE and RBPJKcKO mice KA-injected. **A)** Gene Set Enrichment Analysis (GSEA) using Hallmark gene set from MSigDB Collections highlights three main processes (Inflammatory response, Hypoxia and Apoptosis) affected by the absence of RBPJk, following neuroexcitotoxicity (CRvsKA). GeneMania visualizes the network of interaction of genes within the processes. RNAseq validation using RT-PCR for up- and down-regulated genes in **B)** CRvsKA comparison and **C)** CNvsEE comparison. **D)** Immunoistochemistry of Gabra2α in hippocampal sections of wild-type (wt) mice, RBPJKcKO mice and Notch1cKO mice shows increase of Gabra2 protein level in absence of RBPJk and more marginally in Notch1cKO. *** indicates P values < 0.001, * is for pvalue < 0,05, and indicates p-value <0.07. Scale bar in D is 500 *μ*m

### 3.3. Validation of selected Notch targets

Comparative gene set enrichment analysis (GSEA) of the DEGs in KA vs CC and CR vs KA shows a strong complementarity of the hallmarked pathways of inflammation (IL-2-Stat4 and MAPK-IL-6-JAK-STAT3), hypoxia (TNFA-NFKB) and apoptosis (p53 and G2M) (Figure 3A) confirming that the absence of RBPJK in neurons has an instrumental effects on neuronal plasticity and viability. To confirm the reproducibility of the RNAseq analysis, we have performed RT-PCR (n= 4 for each genotype/condition) validation of selected hits with a 2-fold differential expression change with neuronal (Gabra2 and Bdnf) and non-neuronal identity (Plekha2 and Pttg1). We confirmed a significant downregulation of Bdnf in absence of RBPJK under KA treatment (63%, p=0.03) (Figure 3B), Plekha2 and Pttg1 were comparably slightly repressed in absence of Notch1 (25%, p=0.04 and p=0.01) (Figure 3C), while Gabra2 tends to increase in absence of RBPJK (3.35 Folds, p=0.06) as well as Notch1 (1.5 Folds, p<0.001) (Figure 3B and 3C). Gabra2 upregulation was further confirmed at the protein level by immunohistochemistry of Gabra2 subunit alpha in RBPJKcKO model and upon KA-mediated excitotoxicity, while in Notch1cKO Gabra2a protein levels are not significantly different than wildtype controls (Figure 3D). It was proposed that GABAergic neurotransmission plays a role in the Notch/RBPJK mediated regulation of neuronal plasticity, learning, and memory[59]. In particular, Notch might enhance synaptic transmission and plasticity by reducing the extracellular GABA concentration[60]. This suggests that through canonical signaling, Notch1 might somehow repress the expression of the postsynaptic GABA receptor subunit, affecting Excitatory/ Inhibitory (E/I) transmission.

### 3.4. RBPJK-dependent target genes

To further explore direct Notch target genes in adult neurons, we interrogated each dataset for the occurrence of the RBPJK Consensus Sequence (CSS) CGTGGGAA bound specifically by the transcription factor, RBPJK [61]. RBPJK is typically engaged in a transcriptional repressor complex, that is transformed from a repressor to an activator complex when Notch is activated/cleaved and the intracellular domain (NICD) enters the nucleus. NICD binds to RBPJK, displacing the repressive cofactors bound to RBPJK and recruits a transcriptional activator complex, which initiates transcription of its target genes[62]. We compared the RBPJK CSS occurrence with the one of random motifs commonly found at promoters (n=500). Among up-regulated genes, we found that KA-induced excitotoxicity enriches for genes carrying RBPJK CSS in their promoter. These genes remain low in EE and, as expected for LOF models, RBPJK motif doesn’t appear enriched either in Notch1cKO (CN) nor in RBPJKcKO (CR) (Figure 4A). Intersection of the 132 RBPJK targets found upon KA treatment with the 473 DEGs in RBPJKcKO, identifies 62 genes that represent putative Notch canonical target genes activated upon neuro excitotoxicity (Figure 4B). The heatmap shows that these putative targets have opposite expression’s trends in RBPJKcKO, with the majority of them being downregulated in absence of RBPJK (Figure 4C). Taken together, these results confirm that activation of Notch/RBPJK signaling in neurons upon KA injury exerts a strong transcriptional drive, and that lack of RBPJK blocks the transcriptional program with an overall neuroprotective effect [41, 63]. To validate thein *silico* identified target genes, we combined transcriptome analysis with Chromatin Immunoprecipitation (ChIP) assay using RBPJK, NICD specific antibody and FBS as negative control [54]. The experiment was performed for canonical targets (Hes1 and Hes5) and selected putative targets with known association with AD (Npy, Nptx2, Pdch8, Scg2 and Tnc-C), using Fbw7 as a negative control. Results confirm the direct binding of RBPJK to the promoter region. Precipitation with NICD results less efficiently but shows the same trend (Figure 5A). From the list of direct targets, the strongest bound is Nptx2, an early immediate gene in hippocampal ensembles, with synaptic scaling function and a driver of cognitive decline in AD[64]. Among the putative targets, we selected Nptx2, Npy, Pdch8 and Tnc-C which are implicated in synaptic plasticity and have been functionally associated with Alzheimer’s disease pathophysiology to confirm the dependecies to canonical Notch-RBPJK signaling. We confirmed using RTPCR (n= 4 for each genotype/condition) that in RBPJKcKO, Nptx2 and Tnc-C are significantly reduced while Npy and Pdch8 are only subtly changed (Figure 5B). The loss of Nptx2 in RBPJKcKO was confirmed using fluorescent immunohistochemistry on hippocampal sections [Integrated Density (IntDen) of wt=96.4 ±18.3, IntDen of RBPJKcKO=27.35 ±1.6, p=0.02; Figure 5C], which resulted in a comparable repression of GluR1 (Figure 5D). The present data corroborates the role of Notch as an important driver of synaptic plasticity in neurons with direct associations with AD neuronal networks’ dysfunction.

**Figure 4:**
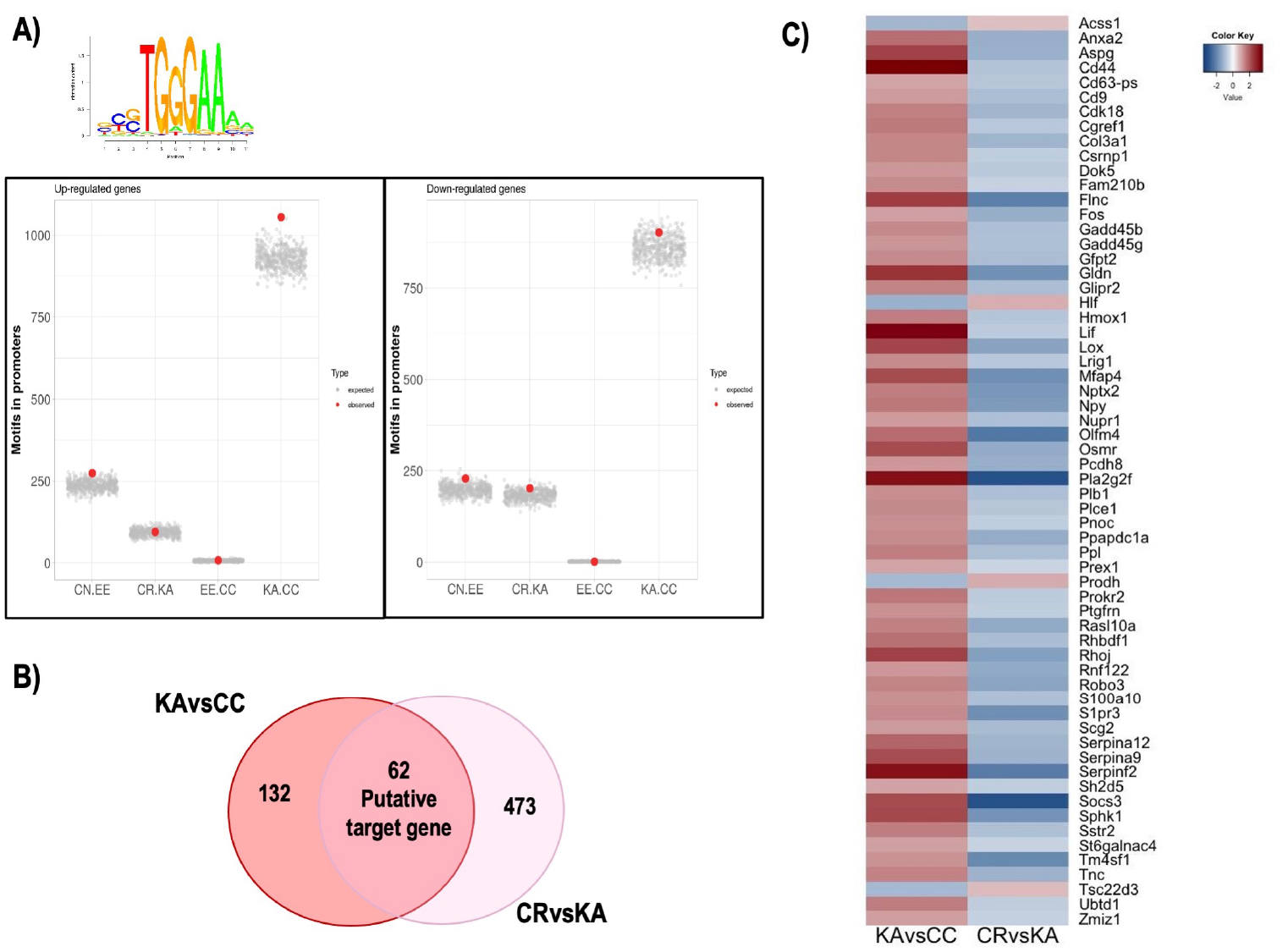
Kainic acid excitotoxicity induces upregulation of RBPJK-dependent genes. **A)** *In silico* analysis of RBPJK consensus sequence on the promoter regions of differential expressed genes in the comparisons: Environmental Enrichment (EE) against Environmental Enrichment in absence of Notch1 (CN), Kainate (KA) versus Kainate in absence of RBPJK (CR), Cage Control (CC) vs Environmental Enrichment (EE), and Kainate (KA) versus Cage Control condition (CC) shows a significative enrichment for upregulated genes upon Kainate treatment (KA.CC). The plot represents the occurrence of RBPJK motif (red dot) compared to random expected promoters (n=500, grey dots). **B)** Matching between the identified genes in the KAvsCC carrying RBPJK CSS, with CRvsKA dataset identifies 62 putative targets. **C)** Heat map shows the specular fold changes in expression between KAvsCC and CRvsKA.

**Figure 5:**
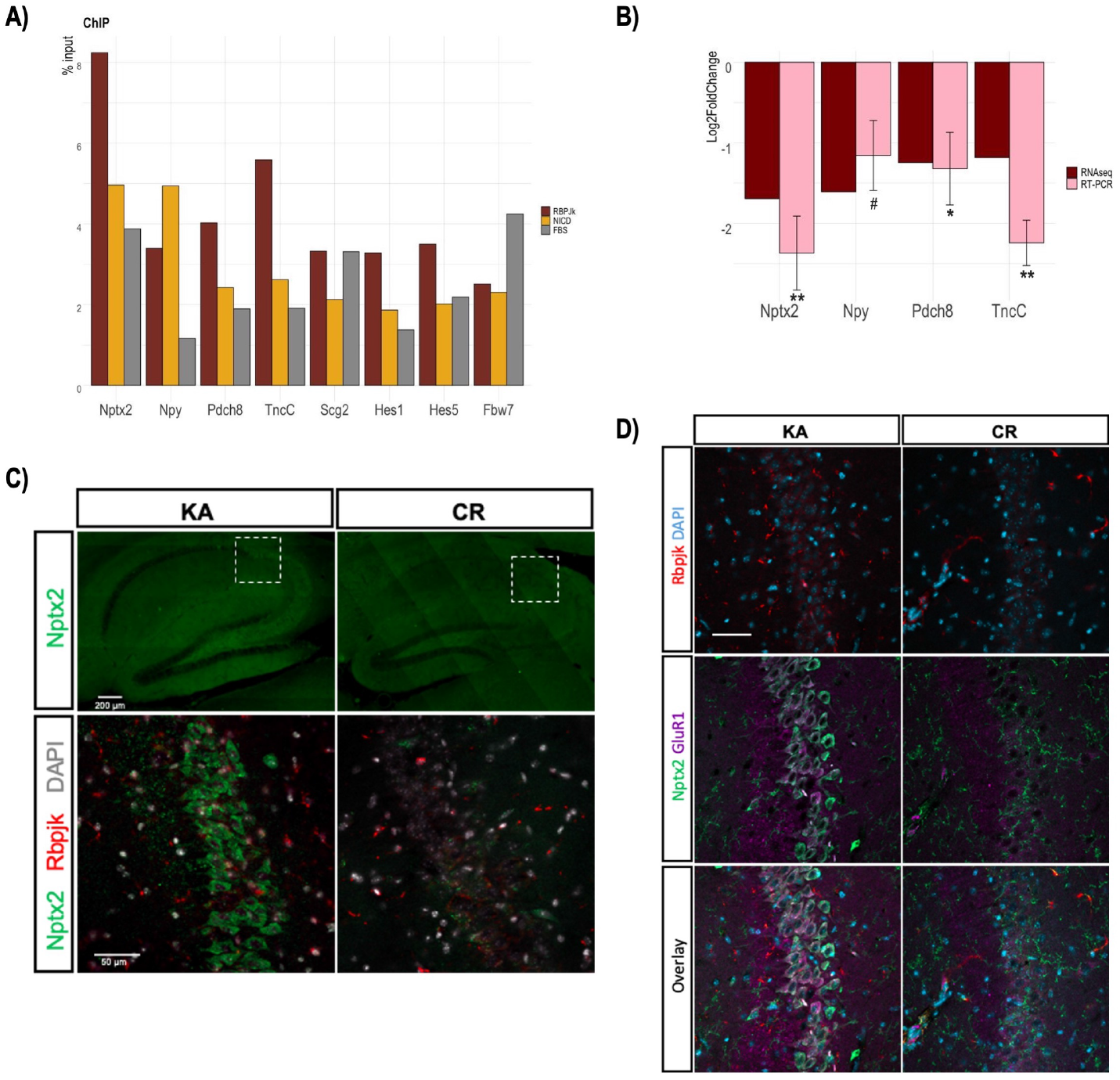
Notch1/RBPJK canonical signaling regulates expression levels of neuroprotective genes upon KA-induced neuroexcitotoxicity. **A)** ChIP-qPCR analysis of Notch1 canonical targets (Hes1 and Hes5) and selected target genes (Nptx2, Npy, Pdch8, Tnc-C and Scg2) confirms the binding of RBPJK and NICD proteins to a subset of canonical and putative target genes. FBS is used as negative control antibody, Fbw7 is the negative control target. **B)** RT-PCR analysis of selected targets with reduced expression (log2FC) in RBPJKcKO after KA treatment. **C)** Double immunolabeling for Rbpjk and Nptx2 shows high expression of Nptx2 protein in hippocampal pyramidal neurons of mice KA-injected (KA). Nptx2 expression is abolished in RBPJKcKO mice (CR). **D)** Representative immunostaining of Rbpjk, and Nptx2 and glutamate receptor 1 (GluR1, AMPA subtype) show concomitant reduction of GluR1 and Nptx2 in RBPJKcKO mice(CR) upon neuroexcitotoxicity. ** indicates pvalue < 0.01, * is for pvalue < 0.05, and indicates p-value < 0.07.

### 3.5. Notch signaling during Alzheimer’s Disease progression

The RNA-seq in the rodent model helped us to computationally identify and validate a list of direct and indirect target genes downstream Notch signaling, with a potential role in the progression of Alzheimer’s Disease dementia. We decided to explore the expression profiles of Notch signaling components and the discovered targets in post-mortem entorhinal cortices from a multicentric cohort of subjects with mild-moderate AD (n=11), severe AD (n=16), and age-matched healthy controls (CTL, n=11). Molecular analyses in humans evidenced a great variability among samples, due to the small number of replicates, and don’t provide cell-type resolution. Nevertheless, it is possible to appreciate a certain trend of Notch signaling expression in the entorhinal cortex during the progression of the disease. We first investigated the expression of some signaling components: transcription of RBPJK increases in both moderate and severe AD pathology compared to healthy controls. In contrast, Notch receptor expression appears unchanged in this dataset. Canonical targets have a more fluctuating expression with Hes1 showing progressive upregulation from moderate to severe AD and Hes5 and Hey1 being more expressed during the moderate stage. Interestingly, Hes5 expression drops drammat-ically in severe AD and its expression correlates with Gabra2a and Map2 while it’s inversely correlated to GFAP, supporting a neuronal expression of this target. The ligand DNER seems to increase throughout the pathology, while Jagged1 doesn’t change much its expression (Figure 6C). These results reflect the trend observed in the RNAseq of KA insulted mice (Figure 2). Correlation analysis reveals that Notch1, RBPJK, Jagged1, DNER and Hes1 and Hey1 expressions (Figure 6F) are significantly correlated confirming the pathway architecture in human brain tissue in the context of AD. While there is general upregulation of Notch signaling with the severity of AD, the expression of the newly discovered target genes (Nptx2, Npy, and Scg2) is reduced during severe AD pathology (Figure 6D). A Similar trend can be observed for the indirect target BDNF that is upregulated during the moderate stage of AD, alongside RBPJK transcription, but it decreases with the progression of the disease (Figure 6E). Similarly, Gabra2a mRNA levels are upregulated during the moderate phase but later in severe AD decrease (Figure 6E). Overall the investigated plasticity genes tend to decrease in the severe AD with Scg2 and Gabra2a being highly correlated (Figure 6G) and supporting a loss of synaptic efficiency and integrity. Expression of Annexin II (Anxa2) is limited to glia and in particular it is found associated to processes of reactive astrocytes. During AD, it is expressed by astrocytes closely associated with beta-amyloid plaques[65]. From our RNAseq, Anxa2 is strongly downregulated, when RBPJK lacks in neurons. RT-PCR of AD patients shows a reduction of Anxa2 in moderate AD, while mRNA level increases in severe AD (Figure 6F). Taken the glial origin of Anxa2, it is possible that it is glial Notch to regulate Anxa2 expression, rather than neuronal Notch. Little is known about Plekha2’s involvement in Alzheimer’s disease but similarly to the plasticity genes, transcriptional downregulation follows the trend of Scg2, Bdnf and Gabra2a in the disease progression (Figure 6F). The present transcriptional profiling confirms that Notch signaling acts in concert in the progression of AD but functional dependencies with genes that are specific to neurons it is at time shadowed based on the pathway’s ubiquitary presence.

**Figure 6:**
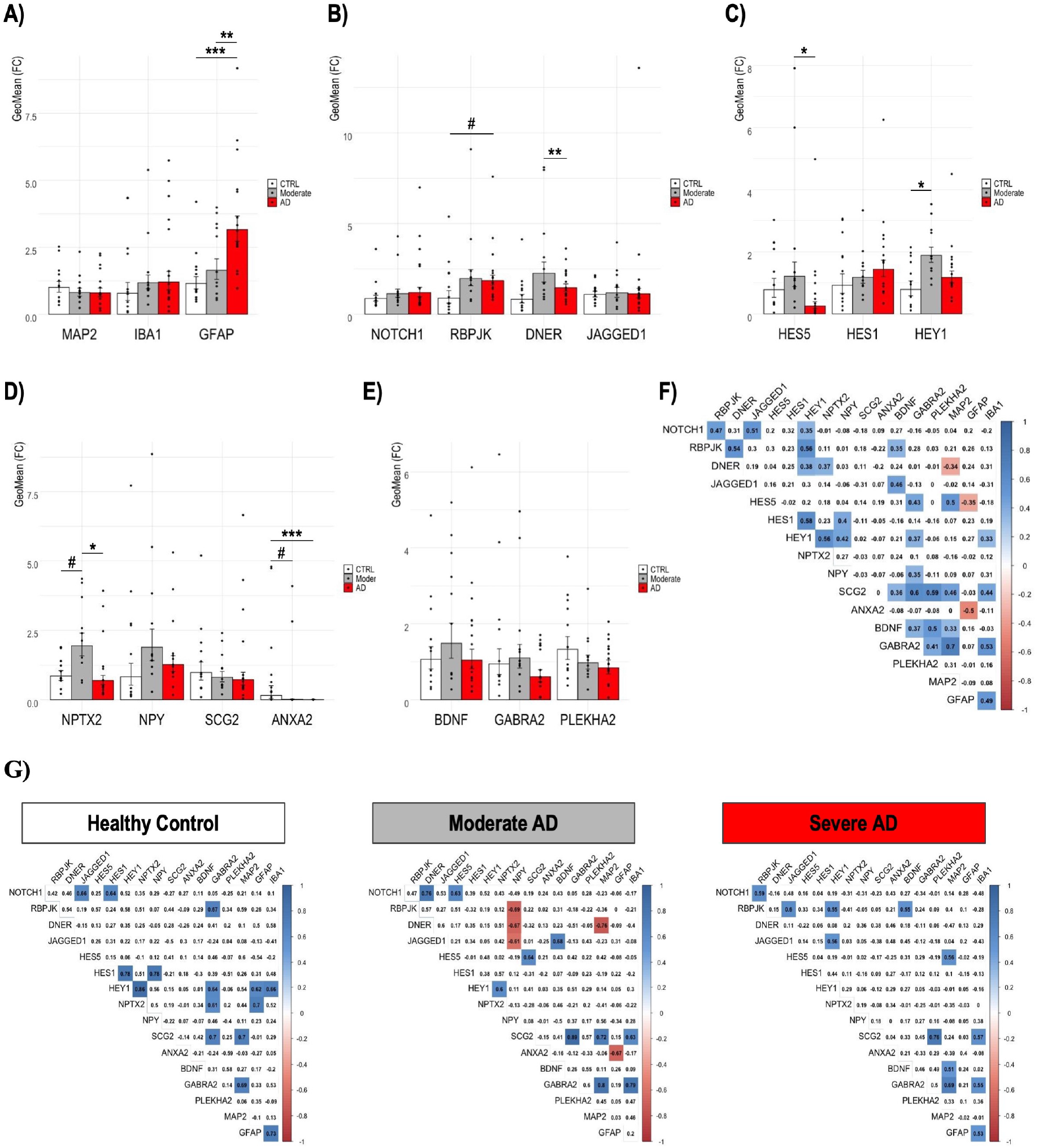
Notch signaling genes and RBPJK-dependent genes are misregulated during Alzheimer’s Disease progression. Bar plots with Jitter show mRNA expression measured with RT-PCR on entorhinal cortex samples containing the hippocampus of Alzheimer’s Disease patients (n=10), AD patients with Moderate disease (n=5) and Control subjects (n=9). **A)** cell type markers, **B)** Notch1 signaling components and **C)** canonical targets, **D)** Notch1 newly discovered direct targets and **E)**Notch1 newly discovered indirect target are analyzed. Spearman’s rank correlation matrix of mRNA expression of Notch signaling genes and neuronal or non-neuronal genes in human entorhinal cortices **F)** in aggregate and **G)** divided by condition. For the bar plot, data are represented as Geometric mean of fold change ± Geometric SEM relative to healthy controls. *** indicates P values < 0.001, ** is for pvalue < 0.01, * is for pvalue < 0,05, and indicates p-value < 0.07

## 4. Discussion

In the adult brain, Notch signaling plays an important role in neuronal stem cell maintenance, morphogenesis, learning and memory processing as well as neuronal loss. The context and the condition in which this cascade plays out determines largely its outcome. Nevertheless, the molecular mechanisms underlying its polyedric functions remain to date largely untapped. This has limited our understanding of whether the loss or the increase in Notch observed in AD brains may be clinically relevant. Notch1 is one of the functional targets of the gamma secretase complex[33]. Presenilin mutations affect Notch activation and its signaling cascade inspiring alternative hypothesis than amyloid-β on the pathogenesis of AD. We have previously shown that Notch1 behaves similarly to an early immediate molecule with induction in expression and signaling upon increased synaptic transmission during spatial memory encoding. However, upon glutamate spillover, in condition of ischemic or epileptic injury, Notch activity is aberrantly induced contributing to neuronal demise. To explain its function, using hippocampal specific loss of function mouse models, we demonstrated that ablation of Notch1 or its cognate ligand Jagged1 affect spatial learning and memory by interfering with NMDAR transmission through a non-transcriptional mechanism[26]. As an additional signaling modality, loss of Rbpjk affects memory formation by increasing the inhibitory tone through GAT2 and BGT1 repression and likely through this mechanism confers neuroprotection following epileptic injury[60] Interestingly, shifted excitation/inhibition balance has been suggested as a major contributor to cognitive decline in Alzheimer’s disease and changes in transcription of the GABAA receptor subunit have been noted. In particular, an increase in the transcripts of the α2 subunit was one of the changes observed in temporal cortices of AD brains, along with increase or decrease of transcription for other subunits[66]. The loss of Notch signaling therefore appears to reduce neuronal excitability on one hand through shunting NMDA transmission and enhancing GABAergic transduction. Although how Notch LOF plays out in the memory deficit can be only explained using an omics discovery approach and neuronal specific mutants as employed in this study.

We have confirmed that the majority of genetic changes influenced by Notch are post-synaptic, supporting the role of this pathway in neuronal homeotasis. In absence of both canonical and non-canonical Notch signaling and upon different treatment conditions, Gabra2a mRNA and protein level increase in hippocampal neurons suggesting an increase in inhibitory drive in absence of Notch, which is aligned with the observed increased inhibition in RbjkcKO mice[60]. Also, Plekha2, a lipid binding proteins, which is positively associated to Epilepsy[67] is downregulated in both Notch1cKO and RbjkcKO mice, further supporting a downscaling of synaptic tone. An additional evidence for a direct effect on synaptic plasticity in RBJK LOF, comes from the reduction in Bdnf expression conserved also in humans, confirming our previous finding[35]

While in AD brains, Notch signaling components, Notch1, Rbpjk, DNER and Jagged1 undergo subtle changes, Hes5 and Hey1 targets show distinctive trends. As both Hes and Hey genes are recognized as classical canonical targets[62], this discrepancy suggests that expression of the genes is cell-type specific, which needs validation, or that activation of Hey and Hes genes undergoes differential transcriptional modulation. Furhermore, our RTPCR analysis on AD brains fails to capture a change in these targets in association with Notch/Rbpjk levels but shows a positive correlation between Gabra2a and Plekha2 in the disease continuum further supporting a mechanistic connection. On the other hand, the lack of interplay between Notch components and the two identified targets can be explained by the absence of spatial resolution of the whole tissue transcript analysis or differential mechanisms across the two species. Whereas, the interconnectivity of RBPJK and Notch signaling component expression is confirmed in humans supporting the evolutionary stability of this pathway and giving robustness to the analysis.

In the search for novel transcriptional targets, we have shown that Kainate treatment displays a rise in CSS binding motifs in the differential expressed genes, emphasizing the notion that canonical signaling is strongly induced upon excitotoxic injury [31]. Of those genes, 62 have inverse directionality in the RbpjkcKO hippocampi, further confirming their identity as legitimate transcriptional Notch/Rbpjk targets. Examination of these selected genes with Allen Brain Map Transcriptomic Explorer using whole cortex and hippocampus 10X Database[68] evidenced that the majority (87%) are neuronal-specific. By norrowing our target validation on 5 genes characterized by known association with AD, we confirm that Nptx2, Npy, Pdch8, and TnC but not Scg-2 are direct transcriptional targets of Notch/Rbpjk. Based on the reported function of Nptx2 in Alzheimer’s disease-related inhibitory circuit dysfunction [64], and the observed role of Notch/Rbpjk in regulating E/I balance, we confirmed that in the hippocampus of RbpjkcKO mice Nptx2 levels are strongly reduced causing also a downregulation of GluR1. This is line with the role of Neuronal pentraxins, NPs, as synaptic organizer which herds the specialization and maturation of excitatory synapses through the recruitment of AMPA receptors, GluR1[69] and GluR4[70]. Besides the evidence that synaptic activity induces the expression of NPs[71], we add that this upregulation is a result of Notch/Rbpjk activation. Nptx2 has received particular attention as levels decline in the progression of Alzheimer’s disease and it represents one of the few differential biomarker for amyloid-β resilience[64]. Our analysis in human AD brains shows a fluctuation in the expression of Nptx2 from a rise in Mild-moderate as compared to cognitively normal controls to a decrease in severe AD. Nptx2 transcript levels associate with the canonical target Hey1, validating its identity as Notch/Rbpjk downstream gene. Interestingly, several of the identified Notch-dependent genes, DNER, Hes5, Hey1, Nptx2, Npy, Bdnf and Gabra2a display an upregulation in the mild-moderate stage of Alzheimer’s Disease. This supports that Notch canonical signaling is active and promotes the expression of neuroprotective proteins, likely in the attempt of mitigating the effect of the E/I imbalance in the intermediate stage of the disease.

While this is the first report detailing a number of novel Notch targets, with instrumental functions for synaptic scaling and plasticity which are lost in the progression of AD, we are completely aware of the limitation of this study based on the bulk approach. This makes single-cell resolution methods in human brain tissue imperative to unlock the cell-specific transcriptional maps associated with Notch activity. This is true also for other ubiquitary pathways such as Insulin that have been evoked in the pathogenesis of AD but have failed to provide functional outcomes instrumental for drug development. Finally, despite the fact that we have used a multicentric clinical cohort of brain specimen, our power remains low due to the limited number of cases. This hampers the ability to make conclusive statements about the Notch/Rbpjk signaling ensemble and appeals for consortium studies capable of enriching the datasets with the focus of shedding light on the driving mechanisms underlying AD pathogenesis.

Concluding, this study demonstrates for the first time that Notch/Rbpjk signaling in neurons has direct and indirect targets instrumental for synaptic plasticity substantiating previous work and supporting a role for this molecular cascade in the progression of AD from E/I imbalance to neuronal demise.

## Sample CRediT author statement

**Amalia Perna** Conceptualization, Methodology, Formal analysis, Investigation, Writing - Original Draft, Visualization **Swananda Marathe**: Conceptualization, Methodology, Investigation **Renè Dreos**: Formal analysis **Laurent Falquet**: Formal analysis **Hatice Akarsu**: Formal analysis **Lavinia Alberi**: Conceptualization, Methodology, Resources, Writing - Review Editing, Supervision, Project administration, Funding acquisition

## Acknowledgments

This research was supported by the Swiss National Foundation (SNF-163470; LA). We thank Valérie Tâche and Eva Martin for technical assistance. We acknowledge the help of the undergraduate students who contributed to this project:

## Abbreviations

AD: Alzheimer’s Disease
NFT: Neurofibrillary Tangles
(Aβ): amyloid-beta, E/I, Excitatory/Inhibitory
LOF: Loss of Function
ChIP: Chromatin Immunoprecipitation
CC: Cage Control
KA: Kainic Acid wild type dataset
CR: Kainic Acid RBPJKcKO dataset
EE: Short Enriched Environment wild type dataset
CN: Short Enriched Environment Notch1cKO dataset
GO: Gene Ontology
GSEA: Gene Set Enriched Analysis
CSS: Consensus Sequence.

---

**Figure S1:**
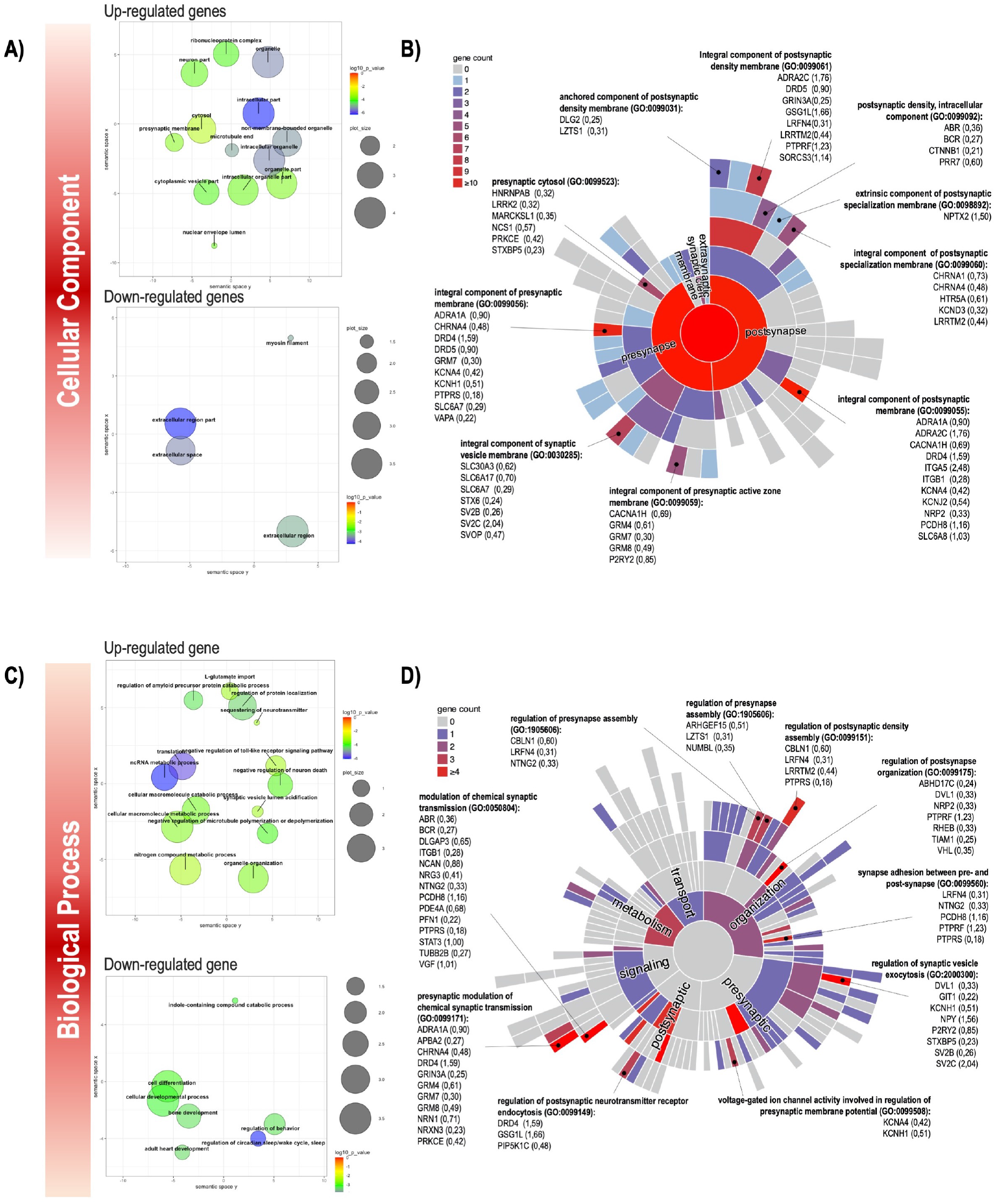
Statistical over-representation analyses of Cellular Component and Biological Process fo KAvsCC dataset. **A)** REViGO Bubbleplot of the Enriched GO Cellular Component cluster representatives of up- and down-regulated genes after KA injection, from GOrilla Enrichment analysis. Circle size is proportional to the frequency of the GO term, whereas color indicates the log10 of p-value for the enrichment derived from GOrilla analysis.**B)** Up-regulated genes visualized using SynGO sunburst plot of gene counts in Cellular Component synaptic terms. Fold change in brackets. **C)** REViGO Bubbleplot of the Enriched GO Cellular Component cluster representatives of up- and down-regulated genes in absence of Notch1 after enriched environment stimulation from Gorilla Enrichment analysis. **D)** SynGO sunburst plot of gene counts in Cellular Component synaptic terms.

**Figure S2:**
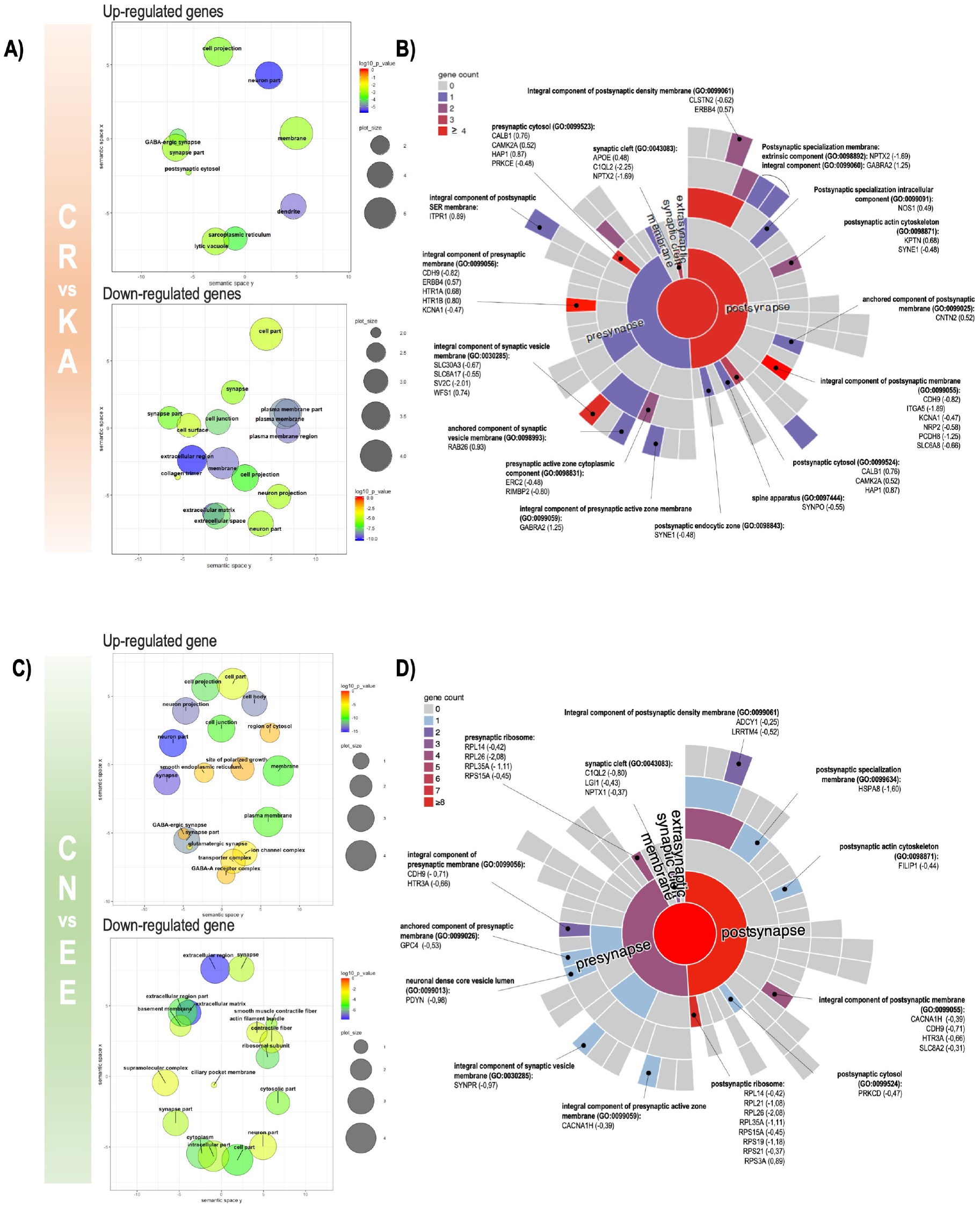
Statistical over-representation analyses of Cellular Component for CRvsKA and CNvsEE datasets. **A)** REViGO Bubbleplot of the Enriched GO Cellular Component cluster representatives of up- and down-regulated genes in absence of RBPJk under neuroexcitotoxicity, from GOrilla Enrichment analysis. Circle size is proportional to the frequency of the GO term, whereas color indicates the log10 of p-value for the enrichment derived from GOrilla analysis.**B)** SynGO sunburst plot of gene counts in Cellular Component synaptic terms for CRvsKA dataset. Fold change in brackets. **C)** REViGO Bubbleplot of the Enriched GO Biological Process cluster representatives of up- and down-regulated genes after KA injection, from Gorilla Enrichment analysis. **D)** SynGO sunburst plot of gene counts in Cellular Component synaptic terms for CNvsEE dataset.

**Figure S3:**
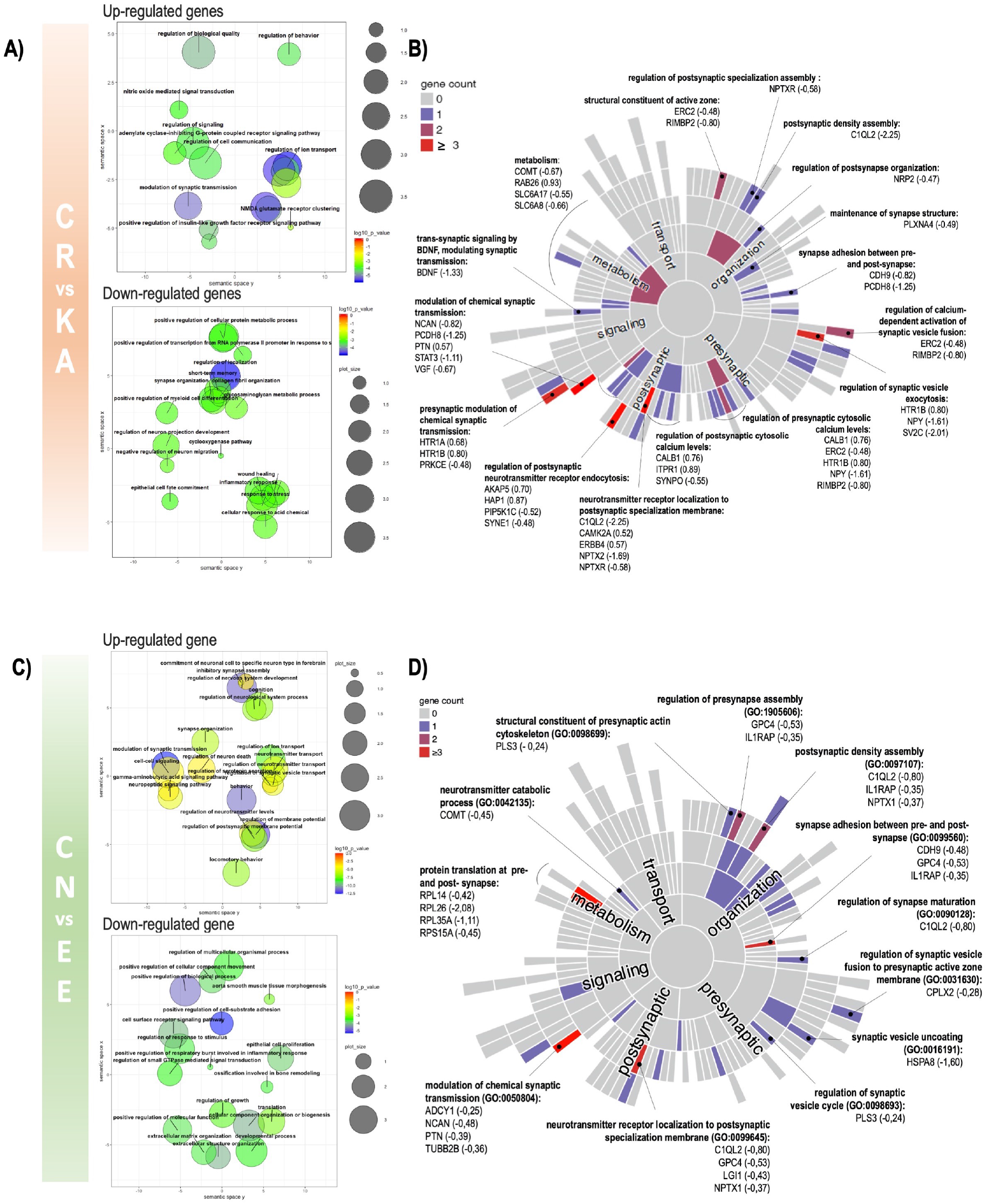
Statistical over-representation analyses of Biological Process for CRvsKA and CNvsEE datasets. **A)** REViGO Bubbleplot of the Enriched GO Biological Process cluster representatives of up- and down-regulated genes in absence of RBPJk under neuroexcitotoxicity, from GOrilla Enrichment analysis. Circle size is proportional to the frequency of the GO term, whereas color indicates the log10 of p-value for the enrichment derived from GOrilla analysis.**B)** SynGO sunburst plot of gene counts in Biological Process synaptic terms for CRvsKA dataset. Fold change in brackets. **C)** REViGO Bubbleplot of the Enriched GO Biological Process cluster representatives of up- and down-regulated genes in absence of Notch1 after enriched environment stimulation from Gorilla Enrichment analysis. **D)** SynGO sunburst plot of gene counts in Biological Process synaptic terms for CNvsEE dataset.

## References

[1] 2020 alzheimer’s disease facts and figures, Alzheimers. Dement.

[2] M. A. DeTure, D. W. Dickson, The neuropathological diagnosis of alzheimer’s disease (2019).

[3] L. M. Osborn, W. Kamphuis, W. J. Wadman, E. M. Hol, Astrogliosis: An integral player in the pathogenesis of alzheimer’s disease, Prog. Neurobiol. 144 (2016) 121–141.

[4] A.-L. Hemonnot, J. Hua, L. Ulmann, H. Hirbec, Microglia in alzheimer disease: Well-Known targets and new opportunities, Front. Aging Neurosci. 11 (2019) 233.

[5] W.-J. Huang, X. Zhang, W.-W. Chen, Role of oxidative stress in alzheimer’s disease, Biomed Rep 4 (5) (2016) 519–522.

[6] W. Wang, F. Zhao, X. Ma, G. Perry, X. Zhu, Mitochondria dysfunction in the pathogenesis of alzheimer’s disease: recent advances (2020).

[7] K. Herrup, The involvement of cell cycle events in the pathogenesis of alzheimer’s disease (2010).

[8] W. J. Jansen, R. Ossenkoppele, D. L. Knol, B. M. Tijms, P. Scheltens, F. R. J. Verhey, P. J. Visser, Amyloid Biomarker Study Group, P. Aalten, D. Aarsland, D. Alcolea, M. Alexander, I. S. Almdahl, S. E. Arnold, I. Baldeiras, H. Barthel, B. N. M. van Berckel, K. Bibeau, K. Blennow, D. J. Brooks, M. A. van Buchem, V. Camus, E. Cavedo, K. Chen, G. Chetelat, A. D. Cohen, A. Drzezga, S. Engelborghs, A. M. Fagan, T. Fladby, A. S. Fleisher, W. M. van der Flier, L. Ford, S. Förster, J. Fortea, N. Foskett, K. S. Frederiksen, Y. Freund-Levi, G. B. Frisoni, L. Froelich, T. Gabryelewicz, K. D. Gill, O. Gkatzima, E. Gómez-Tortosa, M. F. Gordon, T. Grimmer, H. Hampel, L. Hausner, S. Hellwig, S.-K. Herukka, H. Hildebrandt, L. Ishihara, A. Ivanoiu, W. J. Jagust, P. Johannsen, R. Kandimalla, E. Kapaki, A. Klimkowicz-Mrowiec, W. E. Klunk, S. Köhler, N. Koglin, J. Kornhuber, M. G. Kramberger, K. Van Laere, S. M. Landau, D. Y. Lee, M. de Leon, V. Lisetti, A. Lleó, K. Madsen, W. Maier, J. Marcusson, N. Mattsson, A. de Mendonça, O. Meulenbroek, P. T. Meyer, M. A. Mintun, V. Mok, J. L. Molinuevo, H. M. Møllergård, J. C. Morris, B. Mroczko, S. Van der Mussele, D. L. Na, A. Newberg, A. Nordberg, A. Nordlund, G. P. Novak, G. P. Paraskevas, L. Parnetti, G. Perera, O. Peters, J. Popp, S. Prabhakar, G. D. Rabinovici, I. H. G. B. Ramakers, L. Rami, C. Resende de Oliveira, J. O. Rinne, K. M. Rodrigue, E. Rodríguez-Rodríguez, C. M. Roe, U. Rot, C. C. Rowe, E. Rüther, O. Sabri, P. Sanchez-Juan, I. Santana, M. Sarazin, J. Schröder, C. Schütte, S. W. Seo, F. Soetewey, H. Soininen, L. Spiru, H. Struyfs, C. E. Teunissen, M. Tsolaki, R. Vandenberghe, M. M. Verbeek, V. L. Villemagne, S. J. B. Vos, L. J. C. van Waalwijk van Doorn, G. Waldemar, A. Wallin, Å. K. Wallin, J. Wiltfang, D. A. Wolk, M. Zboch, H. Zetterberg, Prevalence of cerebral amyloid pathology in persons without dementia: a meta-analysis, JAMA 313 (19) (2015) 1924–1938.

[9] R. Ossenkoppele, W. J. Jansen, G. D. Rabinovici, D. L. Knol, W. M. van der Flier, B. N. M. van Berckel, P. Scheltens, P. J. Visser, Amyloid PET Study Group, S. C. J. Verfaillie, M. D. Zwan, S. M. Adriaanse, A. A. Lammertsma, F. Barkhof, W. J. Jagust, B. L. Miller, H. J. Rosen, S. M. Landau, V. L. Villemagne, C. C. Rowe, D. Y. Lee, D. L. Na, S. W. Seo, M. Sarazin, C. M. Roe, O. Sabri, H. Barthel, N. Koglin, J. Hodges, C. E. Leyton, R. Vandenberghe, K. van Laere, A. Drzezga, S. Forster, T. Grimmer, P. Sánchez-Juan, J. M. Carril, V. Mok, V. Camus, W. E. Klunk, A. D. Cohen, P. T. Meyer, S. Hellwig, A. Newberg, K. S. Frederiksen, A. S. Fleisher, M. A. Mintun, D. A. Wolk, A. Nordberg, J. O. Rinne, G. Chételat, A. Lleo, R. Blesa, J. Fortea, K. Madsen, K. M. Rodrigue, D. J. Brooks, Prevalence of amyloid PET positivity in dementia syndromes: a metaanalysis, JAMA 313 (19) (2015) 1939–1949.

[10] E. V. Varela, G. Etter, S. Williams, Excitatory-inhibitory imbalance in alzheimer’s disease and therapeutic significance (2019).

[11] A. C. Burggren, G. W. Small, F. W. Sabb, S. Y. Bookheimer, Specificity of brain activation patterns in people at genetic risk for alzheimer disease, Am. J. Geriatr. Psychiatry 10 (1) (2002) 44–51.

[12] M. M. Machulda, D. T. Jones, P. Vemuri, E. McDade, R. Avula, S. Przybelski, B. F. Boeve, D. S. Knopman, R. C. Petersen, C. R. Jack, Jr, Effect of APOE ε4 status on intrinsic network connectivity in cognitively normal elderly subjects, Arch. Neurol. 68 (9) (2011) 1131–1136.

[13] R. A. Sperling, P. S. Laviolette, K. O’Keefe, J. O’Brien, D. M. Rentz, M. Pihlajamaki, G. Marshall, B. T. Hyman, D. J. Selkoe, T. Hedden, R. L. Buckner, J. A. Becker, K. A. Johnson, Amyloid deposition is associated with impaired default network function in older persons without dementia, Neuron 63 (2) (2009) 178–188.

[14] Y. I. Sheline, M. E. Raichle, A. Z. Snyder, J. C. Morris, D. Head, S. Wang, M. A. Mintun, Amyloid plaques disrupt resting state default mode network connectivity in cognitively normal elderly, Biol. Psychiatry 67 (6) (2010) 584–587.

[15] B. C. Dickerson, D. H. Salat, D. N. Greve, E. F. Chua, E. Rand-Giovannetti, D. M. Rentz, L. Bertram, K. Mullin, R. E. Tanzi, D. Blacker, M. S. Albert, R. A. Sperling, Increased hippocampal activation in mild cognitive impairment compared to normal aging and AD (2005).

[16] A. Bakker, G. L. Krauss, M. S. Albert, C. L. Speck, L. R. Jones, C. E. Stark, M. A. Yassa, S. S. Bassett, A. L. Shelton, M. Gallagher, Reduction of hippocampal hyperactivity improves cognition in amnestic mild cognitive impairment, Neuron 74 (3) (2012) 467–474.

[17] R. A. Rissman, W. C. Mobley, Implications for treatment: GABAA receptors in aging, down syndrome and alzheimer’s disease (2011).

[18] M. A. Busche, A. Konnerth, Impairments of neural circuit function in alzheimer’s disease, Philos. Trans. R. Soc. Lond. B Biol. Sci. 371 (1700).

[19] C. Shepherd, H. McCann, G. M. Halliday, Variations in the neuropathology of familial alzheimer’s disease (2009).

[20] N. Brouwers, K. Sleegers, C. Van Broeckhoven, Molecular genetics of alzheimer’s disease: an update, Ann. Med. 40 (8) (2008) 562–583.

[21] R. Potter, B. W. Patterson, D. L. Elbert, V. Ovod, T. Kasten, W. Sigurdson, K. Mawuenyega, T. Blazey, A. Goate, R. Chott, K. E. Yarasheski, D. M. Holtzman, J. C. Morris, T. L. S. Benzinger, R. J. Bateman, Increased in vivo amyloid-42 production, exchange, and loss in presenilin mutation carriers (2013).

[22] O. A. Gomazkov, How do signaling molecules organize higher brain functions? (2015).

[23] K. Gadhave, D. Kumar, V. N. Uversky, R. Giri, A multitude of signaling pathways associated with alzheimer’s disease and their roles in AD pathogenesis and therapy, Med. Res. Rev.

[24] J. Falo-Sanjuan, S. J. Bray, Decoding the notch signal, Dev. Growth Differ. 62 (1) (2020) 4–14.

[25] L. Redmond, S. R. Oh, C. Hicks, G. Weinmaster, A. Ghosh, Nuclear notch1 signaling and the regulation of dendritic development, Nat. Neurosci. 3 (1) (2000) 30–40.

[26] E. Brai, S. Marathe, S. Astori, N. B. Fredj, E. Perry, C. Lamy, A. Scotti, L. Alberi, Notch1 regulates hippocampal plasticity through interaction with the reelin pathway, glutamatergic transmission and CREB signaling, Front. Cell. Neurosci. 9 (2015) 447.

[27] L. Alberi, S. Liu, Y. Wang, R. Badie, C. Smith-Hicks, J. Wu, T. J. Pierfelice, B. Abazyan, M. P. Mattson, D. Kuhl, M. Pletnikov, P. F. Worley, N. Gaiano, Activity-Induced notch signaling in neurons requires Arc/Arg3.1 and is essential for synaptic plasticity in hippocampal networks (2011).

[28] Y. Li, M. A. Hibbs, A. L. Gard, N. A. Shylo, K. Yun, Genome-wide analysis of N1ICD/RBPJ targets in vivo reveals direct transcriptional regulation of wnt, SHH, and hippo pathway effectors by notch1, Stem Cells 30 (4) (2012) 741–752.

[29] K. Yoon, N. Gaiano, Notch signaling in the mammalian central nervous system: insights from mouse mutants, Nat. Neurosci. 8 (6) (2005) 709–715.

[30] P. Andersen, H. Uosaki, L. T. Shenje, C. Kwon, Non-canonical notch signaling: emerging role and mechanism, Trends Cell Biol. 22 (5) (2012) 257–265.

[31] L. Alberi, S. E. Hoey, E. Brai, A. L. Scotti, S. Marathe, Notch signaling in the brain: in good and bad times, Ageing Res. Rev. 12 (3) (2013) 801–814.

[32] T. Moehlmann, E. Winkler, X. Xia, D. Edbauer, J. Murrell, A. Capell, C. Kaether, H. Zheng, B. Ghetti, C. Haass, H. Steiner, Presenilin-1 mutations of leucine 166 equally affect the generation of the notch and APP intracellular domains independent of their effect on abeta 42 production, Proc. Natl. Acad. Sci. U. S. A. 99 (12) (2002) 8025–8030.

[33] M. Okochi, Presenilins mediate a dual intramembranous gamma-secretase cleavage of notch-1, EMBO J. 21 (20) (2002) 5408–5416.

[34] O. Berezovska, M. Q. Xia, B. T. Hyman, Notch is expressed in adult brain, is coexpressed with presenilin-1, and is altered in alzheimer disease, J. Neuropathol. Exp. Neurol. 57 (8) (1998) 738–745.

[35] E. Brai, N. A. Raio, L. Alberi, Notch1 hallmarks fibrillary depositions in sporadic alzheimer’s disease (2016).

[36] T V. Arumugam, S.-H. Baik, P. Balaganapathy, C. G. Sobey, M. P. Mattson, D.-G. Jo, Notch signaling and neuronal death in stroke, Progress in neurobiology 165 (2018) 103–116.

[37] X.-Y. Zheng, H.-L. Zhang, Q. Luo, J. Zhu, Kainic acid-induced neurodegenerative model: potentials and limitations, J. Biomed. Biotechnol. 2011 (2011) 457079.

[38] H. Han, K. Tanigaki, N. Yamamoto, K. Kuroda, M. Yoshimoto, T. Nakahata, K. Ikuta, T. Honjo, Inducible gene knockout of transcription factor recombination signal binding protein-j reveals its essential role in T versus B lineage decision (2002).

[39] J. Z. Tsien, D. F. Chen, D. Gerber, C. Tom, E. H. Mercer, D. J. Anderson, M. Mayford, E. R. Kandel, S. Tonegawa, Subregion- and cell type-restricted gene knockout in mouse brain, Cell 87 (7) (1996) 1317–1326.

[40] F. Radtke, I. Ferrero, A. Wilson, R. Lees, M. Aguet, H. Robson MacDonald, Notch1 deficiency dissociates the intrathymic development of dendritic cells and T cells (2000).

[41] S. Marathe, S. Liu, E. Brai, M. Kaczarowski, L. Alberi, Notch signaling in response to excitotoxicity induces neurodegeneration via erroneous cell cycle reentry, Cell Death Differ. 22 (11) (2015) 1775–1784.

[42] A. Buschler, D. Manahan-Vaughan, Brief environmental enrichment elicits metaplasticity of hippocampal synaptic potentiation in vivo, Front. Behav. Neurosci. 6 (2012) 85.

[43] J. D. Storey, R. Tibshirani, Statistical significance for genomewide studies, Proc. Natl. Acad. Sci. U. S. A. 100 (16) (2003) 9440–9445.

[44] I. V. Kulakovskiy, I. E. Vorontsov, I. S. Yevshin, R. N. Sharipov, A. D. Fedorova, E. I. Rumynskiy, Y. A. Medvedeva, A. Magana-Mora, V. B. Bajic, D. A. Papatsenko, F. A. Kolpakov, V. J. Makeev, HOCOMOCO: towards a complete collection of transcription factor binding models for human and mouse via large-scale ChIP-Seq analysis, Nucleic Acids Res. 46 (D1) (2018) D252–D259.

[45] G. Ambrosini, R. Groux, P. Bucher, PWMScan: a fast tool for scanning entire genomes with a position-specific weight matrix, Bioinformatics 34 (14) (2018) 2483–2484.

[46] M. Lawrence, R. Gentleman, V. Carey, rtracklayer: an R package for interfacing with genome browsers (2009).

[47] E. Eden, R. Navon, I. Steinfeld, D. Lipson, Z. Yakhini, GOrilla: a tool for discovery and visualization of enriched GO terms in ranked gene lists, BMC Bioinformatics 10 (2009) 48.

[48] F. Supek, M. Bošnjak, N. Škunca, T. Šmuc, REVIGO summarizes and visualizes long lists of gene ontology terms, PLoS One 6 (7) (2011) e21800.

[49] A. Schlicker, F. S. Domingues, J. Rahnenführer, T. Lengauer, A new measure for functional similarity of gene products based on gene ontology, BMC Bioinformatics 7 (2006) 302.

[50] F. Koopmans, P. van Nierop, M. Andres-Alonso, A. Byrnes, T. Cijsouw, M. P. Coba, L. N. Cornelisse, R. J. Farrell, H. L. Goldschmidt, D. P. Howrigan, N. K. Hussain, C. Imig, A. P. H. de Jong, H. Jung, M. Kohansalnodehi, B. Kramarz, N. Lipstein, R. C. Lovering, H. MacGillavry, V. Mariano, H. Mi, M. Ninov, D. Osumi-Sutherland, R. Pielot, K.-H. Smalla, H. Tang, K. Tashman, R. F. G. Toonen, C. Verpelli, R. Reig-Viader, K. Watanabe, J. van Weering, T. Achsel, G. Ashrafi, N. Asi, T. C. Brown, P. De Camilli, M. Feuermann, R. E. Foulger, P. Gaudet, A. Joglekar, A. Kanellopoulos, R. Malenka, R. A. Nicoll, C. Pulido, J. de Juan-Sanz, M. Sheng, T. C. Südhof, H. U. Tilgner, C. Bagni, À. Bayés, T. Biederer, N. Brose, J. J. E. Chua, D. C. Dieterich, E. D. Gundelfinger, C. Hoogenraad, R. L. Huganir, R. Jahn, P. S. Kaeser, E. Kim, M. R. Kreutz, P. S. McPherson, B. M. Neale, V. O’Connor, D. Posthuma, T. A. Ryan, C. Sala, G. Feng, S. E. Hyman, P. D. Thomas, A. B. Smit, M. Verhage, SynGO: An Evidence-Based, Expert-Curated knowledge base for the synapse, Neuron 103 (2) (2019) 217–234.e4.

[51] A. Subramanian, P. Tamayo, V. K. Mootha, S. Mukherjee, B. L. Ebert, M. A. Gillette, A. Paulovich, S. L. Pomeroy, T. R. Golub, E. S. Lander, J. P. Mesirov, Gene set enrichment analysis: a knowledge-based approach for interpreting genome-wide expression profiles, Proc. Natl. Acad. Sci. U. S. A. 102 (43) (2005) 15545–15550.

[52] D. Merico, R. Isserlin, O. Stueker, A. Emili, G. D. Bader, Enrichment map: a network-based method for gene-set enrichment visualization and interpretation, PLoS One 5 (11) (2010) e13984.

[53] J. Montojo, K. Zuberi, H. Rodriguez, F. Kazi, G. Wright, S. L. Donaldson, Q. Morris, G. D. Bader, GeneMANIA cytoscape plugin: fast gene function predictions on the desktop (2010).

[54] A. Perna, L. A. Alberi, TF-ChIP method for Tissue-Specific gene targets, Front. Cell. Neurosci. 13 (2019) 95.

[55] S. K. Mahata, J. Marksteiner, G. Sperk, M. Mahata, B. Gruber, R. Fischer-Colbrie, H. Winkler, Temporal lobe epilepsy of the rat: differential expression of mRNAs of chromogranin b, secretogranin II, synaptin/synaptophysin and p65 in subfields of the hippocampus (1992).

[56] E.-J. D. Lin, D. Young, K. Baer, H. Herzog, M. J. During, Differential actions of NPY on seizure modulation via Y1 and Y2 receptors: evidence from receptor knockout mice, Epilepsia 47 (4) (2006) 773–780.

[57] N. Otani, H. Nawashiro, N. Nomura, S. Fukui, N. Tsuzuki, S. Ishihara, K. Shima, A role of glial fibrillary acidic protein in hippocampal degeneration after cerebral trauma or kainate-induced seizure (2003).

[58] S. B. Bausch, Potential roles for hyaluronan and CD44 in kainic acid-induced mossy fiber sprouting in organotypic hippocampal slice cultures, Neuroscience 143 (1) (2006) 339–350.

[59] G. A. Hortopan, M. T. Dinday, S. C. Baraban, Spontaneous seizures and altered gene expression in GABA signaling pathways in a mind bomb mutant zebrafish (2010).

[60] S. Liu, Y. Wang, P. F. Worley, M. P. Mattson, N. Gaiano, The canonical notch pathway effector RBP-J regulates neuronal plasticity and expression of GABA transporters in hippocampal networks, Hippocampus 25 (5) (2015) 670–678.

[61] T. Tun, Y. Hamaguchi, N. Matsunami, T. Furukawa, T. Honjo, M. Kawaichi, Recognition sequence of a highly conserved DNA binding protein RBP-Jx (1994).

[62] R. Kopan, M. X. G. Ilagan, The canonical notch signaling pathway: unfolding the activation mechanism, Cell 137 (2) (2009) 216–233.

[63] T. V. Arumugam, Y.-L. Cheng, Y. Choi, Y.-H. Choi, S. Yang, Y.-K. Yun, J.-S. Park, D. K. Yang, J. Thundyil, M. Gelderblom, Others, Evidence that γ-secretase-mediated notch signaling induces neuronal cell death via the nuclear factor-κB-Bcl-2-interacting mediator of cell death pathway in ischemic stroke, Mol. Pharmacol. 80 (1) (2011) 23–31.

[64] M.-F. Xiao, D. Xu, M. T. Craig, K. A. Pelkey, C.-C. Chien, Y. Shi, J. Zhang, S. Resnick, O. Pletnikova, D. Salmon, J. Brewer, S. Edland, J. Wegiel, B. Tycko, A. Savonenko, R. H. Reeves, J. C. Troncoso, C. J. McBain, D. Galasko, P. F. Worley, NPTX2 and cognitive dysfunction in alzheimer’s disease, Elife 6.

[65] D. A. Eberhard, M. D. Brown, S. R. VandenBerg, Alterations of annexin expression in pathological neuronal and glial reactions. immunohistochemical localization of annexins i, II (p36 and p11 subunits), IV, and VI in the human hippocampus, Am. J. Pathol. 145 (3) (1994) 640–649.

[66] A. Limon, J. M. Reyes-Ruiz, R. Miledi, Loss of functional GABAA receptors in the alzheimer diseased brain (2012).

[67] R. S. Sprissler, J. L. Wagnon, R. K. Bunton-Stasyshyn, M. H. Meisler, M. F. Hammer, Altered gene expression profile in a mouse model of scn8a encephalopathy, Experimental neurology 288 (2017) 134–141.

[68] Z. Yao, T. N. Nguyen, C. T. van Velthoven, J. Goldy, A. E. Sedeno-Cortes, F. Baftizadeh, D. Bertagnolli, T. Casper, K. Crichton, S.-L. Ding, et al., A taxonomy of transcriptomic cell types across the isocortex and hippocampal formation.

[69] S.-J. Lee, M. Wei, C. Zhang, S. Maxeiner, C. Pak, S. C. Botelho, J. Trotter, F. H. Sterky, T. C. Südhof, Presynaptic neuronal pentraxin receptor organizes excitatory and inhibitory synapses, Journal of Neuroscience 37 (5) (2017) 1062–1080.

[70] G.-M. Sia, J.-C. Béïque, G. Rumbaugh, R. Cho, P. F. Worley, R. L. Huganir, Interaction of the n-terminal domain of the ampa receptor glur4 subunit with the neuronal pentraxin np1 mediates glur4 synaptic recruitment, Neuron 55 (1) (2007) 87–102.

[71] J. H. Leslie, E. Nedivi, Activity-regulated genes as mediators of neural circuit plasticity, Progress in neurobiology 94 (3) (2011) 223–237.

