## Supplementary figures and images for "Revealing Notch-dependencies in synaptic targets associated with Alzheimer’s disease"

### Supplementary Figure 1

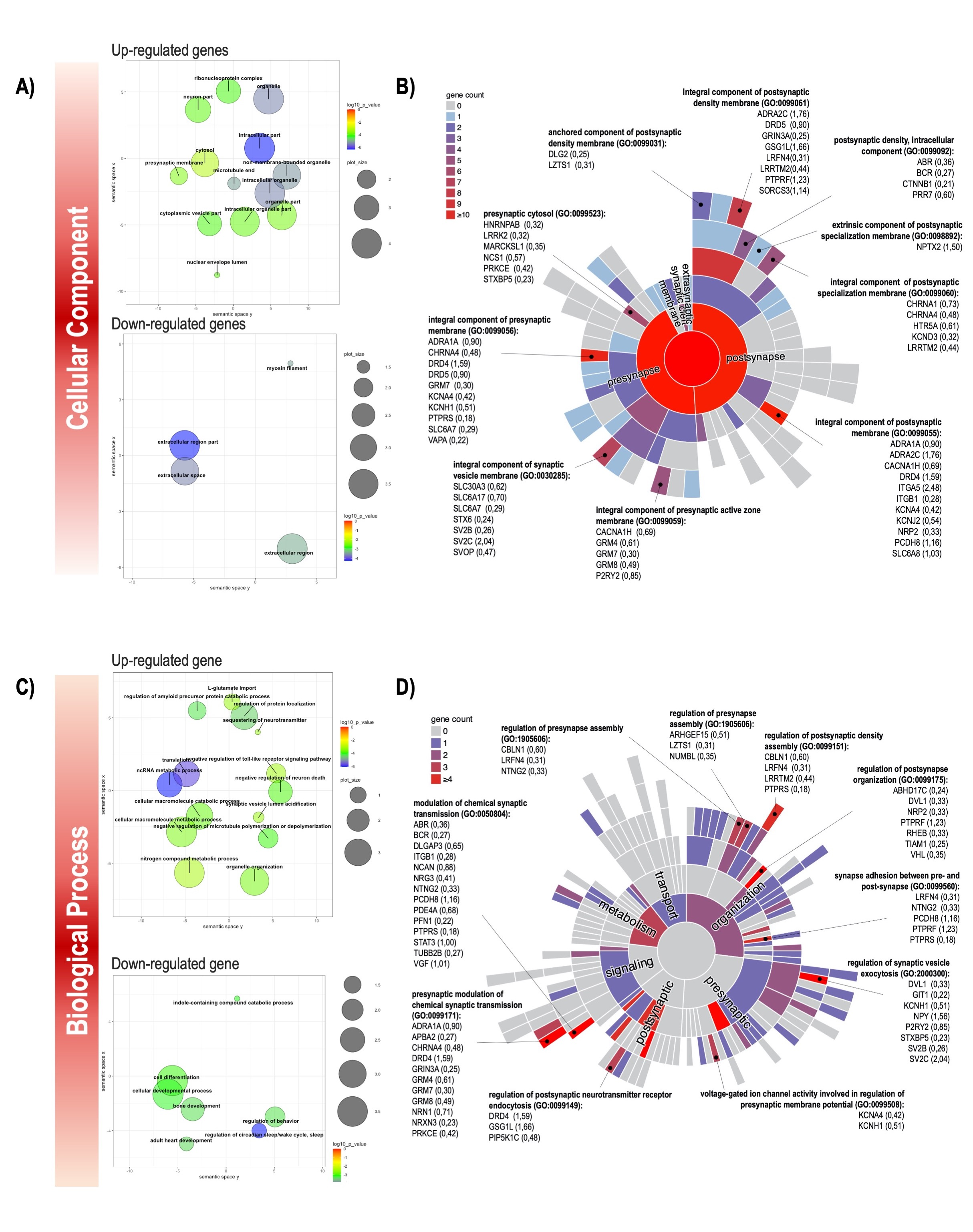

### Supplementary Figure 2

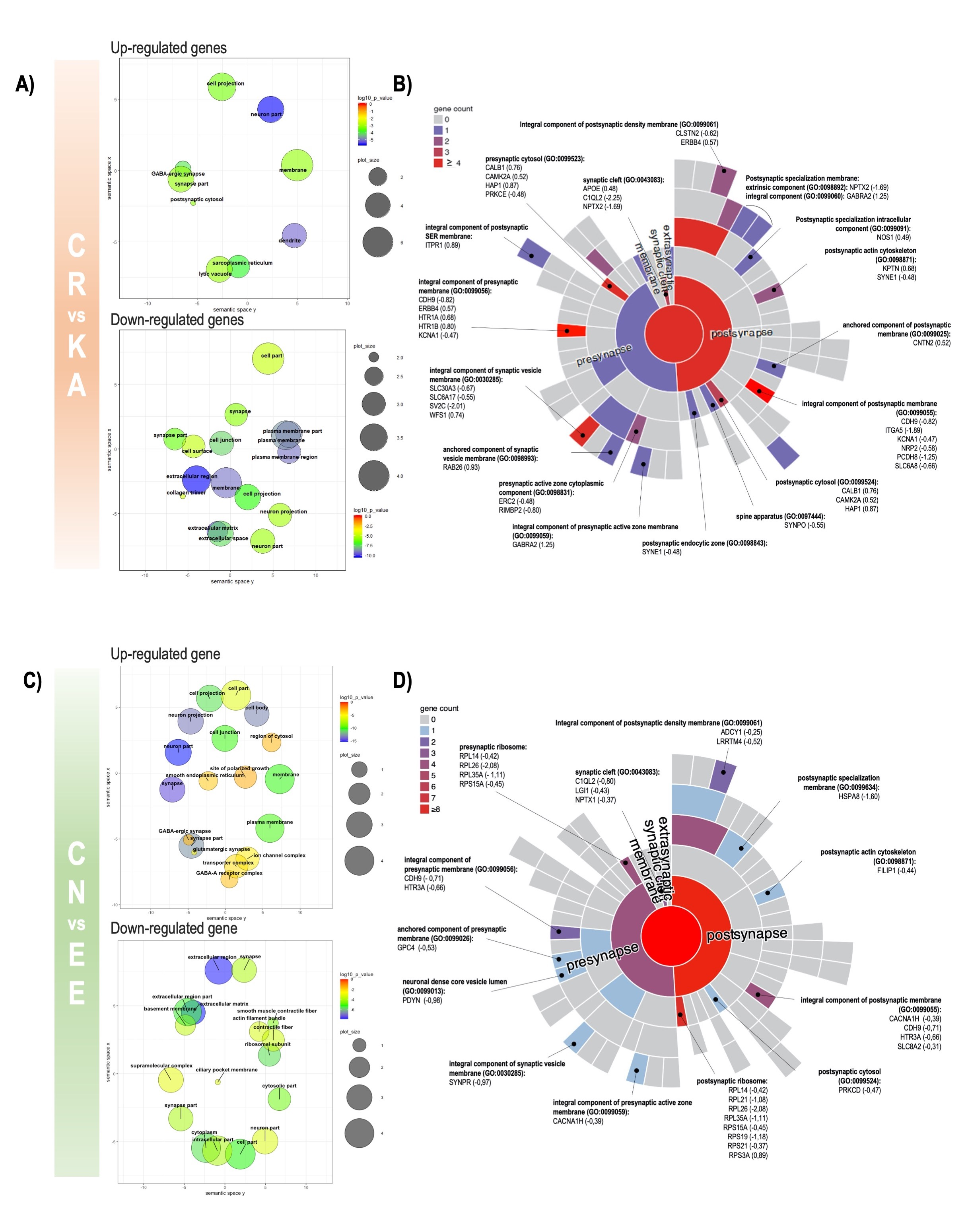

### Supplementary Figure 3

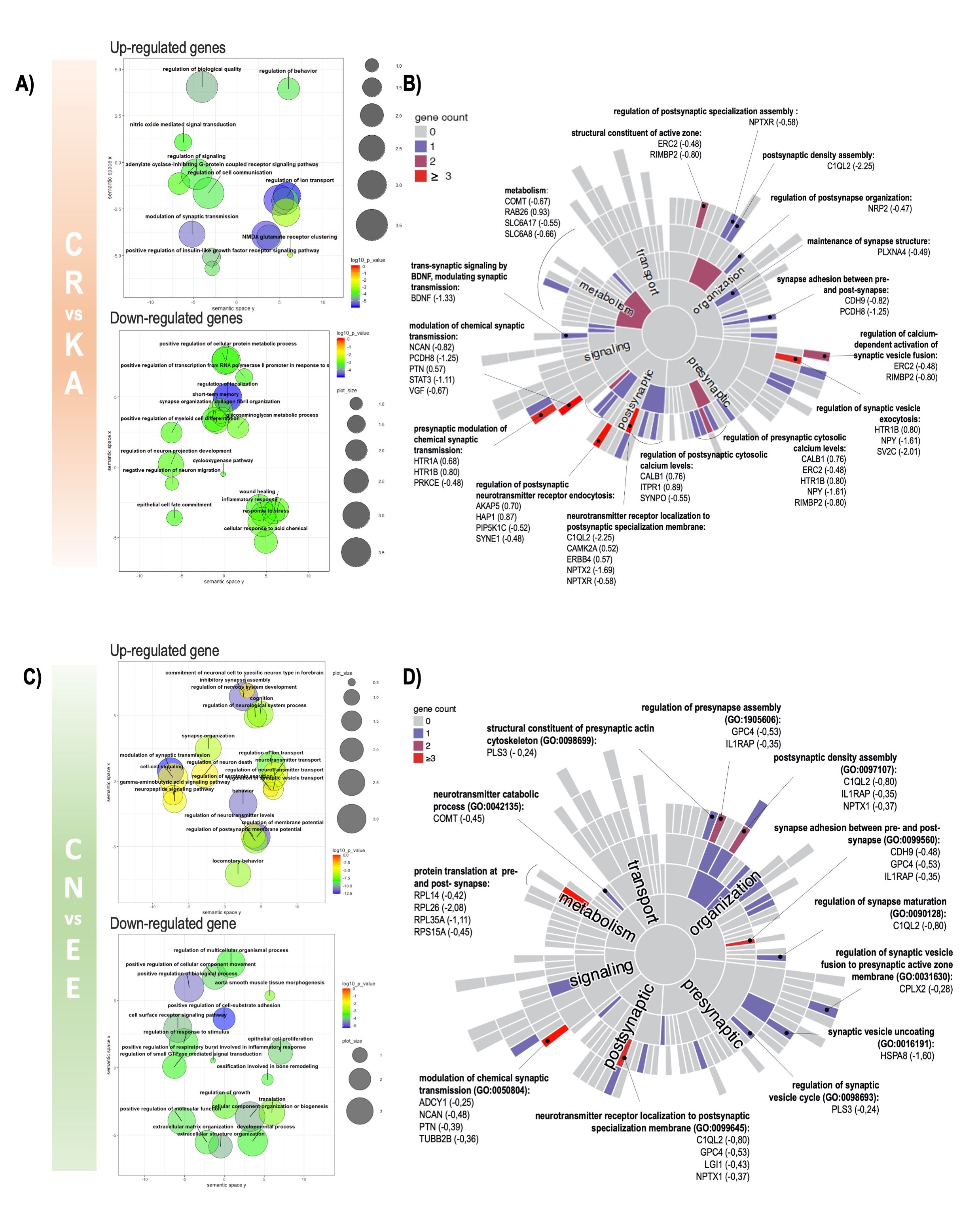
